# Metabolism of bilirubin to urobilinogen by human gut bacteria occurs by a branched, multi-enzyme pathway

**DOI:** 10.64898/2026.06.10.731317

**Authors:** Baylee J. Russell, Erik J. Hasenoehrl, Victoria M. Marando, Joy Lu, Nicholas H. Dang, Davis H. Smith, Kelsey N. Thompson, Jessica M. Chen, Michael J. James, Mudita Goyal, Suzanne Walker, Curtis Huttenhower, Seth Rakoff-Nahoum, Marco Jost

## Abstract

Bilirubin, the predominant product of heme catabolism in mammals, enters the intestine via the hepatobiliary system and subsequently is metabolized by the gut microbiome into multiple downstream derivatives, such as urobilinogen and stercobilinogen. Variation in levels of bilirubin and its derivatives are associated with inflammatory and metabolic disorders, but the microbial species and enzymes that metabolize bilirubin have remained largely unknown, which has precluded causal and mechanistic studies. Here, we demonstrate that metabolism of bilirubin to urobilinogen requires multiple distinct reactions carried out by separate enzymes. We find that bilirubin reductase (BilR), an enzyme that was recently discovered and proposed to convert bilirubin to urobilinogen, is specific for reducing the methine bridges of bilinoids, converting bilirubin to the previously uncharacterized intermediate divinylurobilinogen and mesobilirubin to urobilinogen. We identify bilinoid vinyl reductase (BilV), a flavin-dependent oxidoreductase of the Old Yellow Enzyme (OYE) superfamily, as the enzyme responsible for reducing the vinyl groups of bilirubin and divinylurobilinogen. Both BilR and BilV are capable of acting on multiple bilinoids, such that the reductions required to convert bilirubin to urobilinogen can proceed in multiple orders through a branched network of intermediates. BilV is widespread in human gut metagenomes and is broadly distributed across gut bacteria in a manner that overlaps with but does not mirror the distribution of BilR. These findings establish the identities of novel factors in bacterial bilirubin metabolism, delineate the branched architecture of the pathway, and enable defined studies to interrogate how such branched metabolism contributes to human health and disease.

## Introduction

Gut bacterial transformations of human-derived metabolites play critical roles in human health by altering levels of endogenous ligands for host receptors and generating new bioactive compounds. Key examples include transformations of bile acids and steroids, which occur through complex, branching pathways and the products of which are implicated in effects on T cell polarization, anti-tumor immunity, and central metabolism, among others^1–5^. The broader scope and mechanisms of gut bacterial conversions of host-derived metabolites across the diverse landscape of potential substrates remain largely unknown.

A major host-derived substrate accessible to gut bacteria is bilirubin, the predominant product of heme catabolism in mammals. Bilirubin is generated through oxidative cleavage of heme to form biliverdin and subsequent reduction to bilirubin^6^ and eliminated by conjugation to glucuronic acid in the liver, excretion into bile, and release into the intestinal lumen. In humans, turnover of red blood cells results in excretion of hundreds of milligrams of bilirubin per day, which are nearly quantitatively metabolized by gut bacteria (**Figure 1A**). Characterized steps include deconjugation of bilirubin glucuronide^7^ followed by progressive reduction of six double bonds in bilirubin: two in vinyl side chains, two in methine bridges, and two in the terminal pyrrole rings (**Figure 1A**). These reactions result in a family of tetrapyrrole metabolites that includes urobilinogen and stercobilinogen, often detected in urine and/or stool as their oxidized counterparts, urobilin and stercobilin^8,9^.

**Figure 1:**
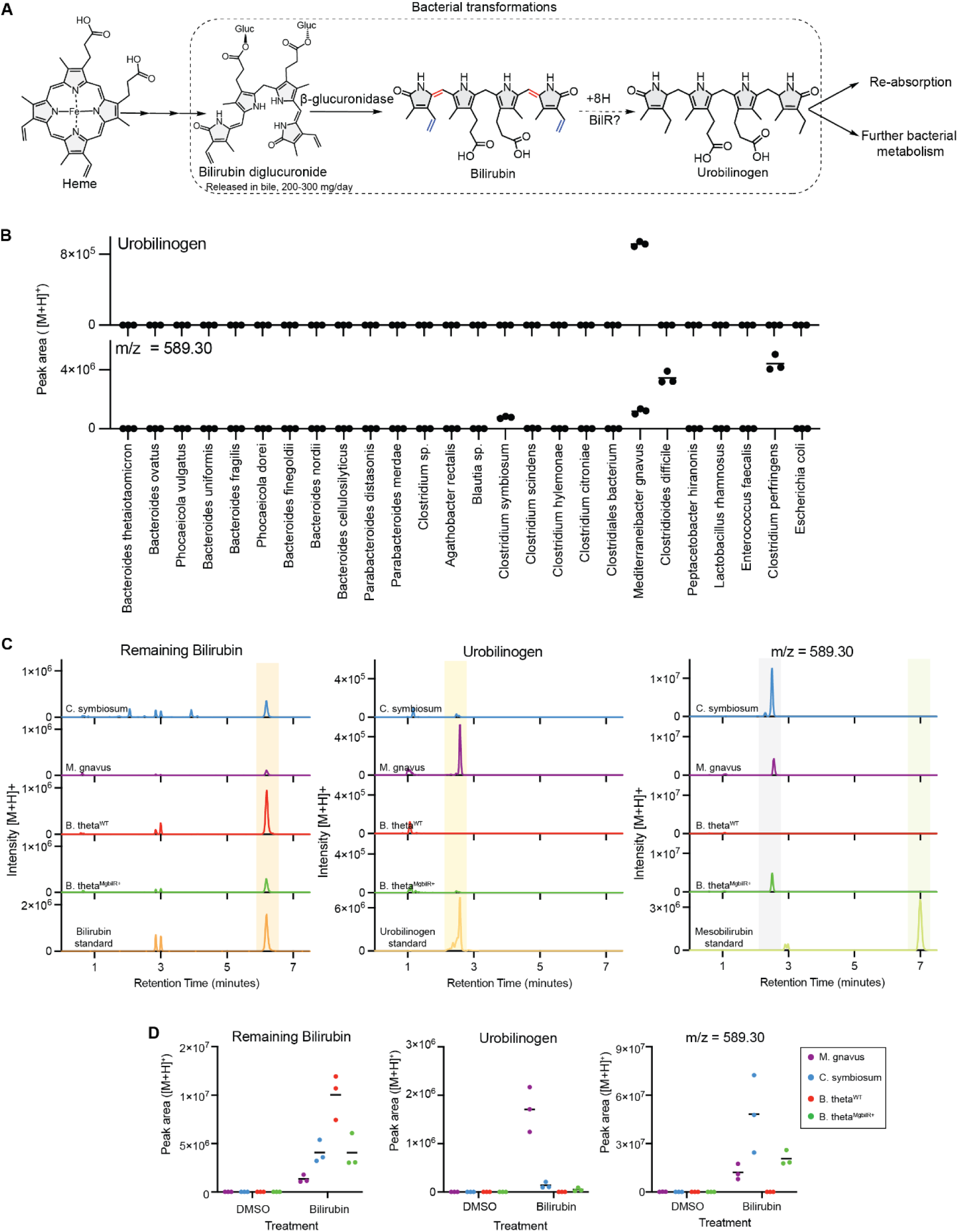
BilR is insufficient to convert bilirubin to urobilinogen. **A.** Overview of bacterial bilirubin transformations. Gluc is glucuronic acid. **B.** Top: Quantified peak area of urobilinogen (standard-verified) from indicated gut bacteria incubated with 10 μM bilirubin for 24 hours. Bottom: Quantified peak area of novel compound with m/z=589.30 and retention time=2.5 min from same cultures. Each dot represents an individual culture. **C.** Representative extracted ion chromatograms (EICs) for bilirubin (left), urobilinogen (middle), or m/z=589.30 (right) after incubation of indicated strains with 10 μM bilirubin for 24 hours. EICs for commercially available standards shown at the bottom. The shaded regions indicate the approximate retention times of the standards (bilirubin: orange; urobilinogen: yellow; mesobilirubin: green) or of the novel compound with m/z=589.30 (gray). **D.** Quantified peak area for bilirubin (left), urobilinogen (middle), or compound with m/z=589.30 and retention time=2.5 min (right). Bilirubin and urobilinogen were verified using standards. Data in B and D are represented as mean and individual values from replicate cultures.

Variation in levels of bilirubin and bacterial bilirubin metabolites have been linked to multiple diseases. Bilirubin was originally considered to be detrimental at elevated levels, such as in kernicterus in neonatal jaundice. More recent studies, however, have associated low systemic bilirubin levels with increased incidence of metabolic syndrome and cardiovascular disease^10–14^, attributed in part to anti-inflammatory and PPARα agonist activities of bilirubin. Conversely, elevated levels of urobilin or urobilinogen are associated with cardiovascular disease and metabolic syndrome in some patient cohorts and in rodent models of disease^14–22^, and suppression of bacterial production of urobilin or urobilinogen is under evaluation as a therapeutic modality^13^. Many of these associations have not resolved urobilin, urobilinogen, or structural isomers and stereoisomers, and some associations diverge for urobilinogen and urobilin^20^, suggesting the possibility of compound-specific effects. Finally, elevated levels of fecal bilirubin and decreased bacterial bilirubin metabolism are associated with increased incidence of inflammatory bowel disease (IBD)^23–26^. Whether these associations reflect causal effects of bacterial consumption of bilirubin and/or activities of the produced metabolites remains unclear, and no biological activities of urobilinogen, urobilin, or other bilirubin metabolites have been defined.

A central obstacle to establishing causal relationships is the lack of knowledge of the bacterial enzymes and species involved in bilirubin metabolism. Although the role of gut bacteria in metabolizing bilirubin was first described over a century ago^8,9^, the only enzyme characterized to date is bilirubin reductase (BilR), a flavin-dependent oxidoreductase encoded by gut Clostridia and Bacilli and proposed to convert bilirubin to urobilinogen^23^. The enzymes that mediate additional steps in bilirubin metabolism remain unknown. Further, recent untargeted mass spectrometry (MS) analyses have identified multiple novel candidate bilirubin metabolites, several of which are depleted in IBD^24^, suggesting a broader landscape of bilirubin metabolites.

Here we sought to define the capacity of human gut bacteria to metabolize bilirubin and the enzymatic basis for metabolism. By screening a taxonomically diverse panel of gut bacteria for metabolism of bilirubin, we find that metabolism of bilirubin to urobilinogen occurs via a multi-step process and requires two reductases with distinct specificities for reducing double bonds in bilirubin. We find that BilR, which was proposed to convert bilirubin to urobilinogen by reducing the vinyl side chains and methine bridges^23^, only reduces the methine bridges and converts bilirubin to divinylurobilinogen, a previously uncharacterized intermediate. Conversely, we identify a novel reductase, bilinoid vinyl reductase (BilV), that is prevalent in the gut microbiome and is responsible for reducing the vinyl groups in bilinoids, including divinylurobilinogen and bilirubin.

BilV was also identified through a genomic approach in a concurrent study^27^. These two enzymes constitute a branched pathway for metabolism of bilirubin. Finally, we performed a bioinformatic survey to create a phylogenetic map of gut bacterial bilirubin metabolism, which revealed overlapping but non-identical distributions of BilV and BilR. Our findings establish the pathway for bacterial metabolism of bilirubin to urobilinogen, identify novel bacterial species that participate in bilirubin metabolism, and enable well-defined studies to interrogate the roles of bilirubin metabolism in human physiology.

## Results

### Reduction of bilirubin to urobilinogen is a multi-enzyme process

To explore bilirubin metabolism by gut bacteria, we assembled a panel of 26 cultured, genome-resolved human gut bacteria that is both taxonomically representative and sufficiently focused for experimental tractability (**Supplemental Table 1**), similar to previous studies^28^. The panel includes multiple strains that encode BilR as well as strains not previously implicated in bilirubin metabolism. We incubated each strain with bilirubin under anaerobic conditions and monitored consumption of bilirubin and appearance of novel products by liquid chromatography– mass spectrometry (LC–MS). We first inspected the data for the transformation of bilirubin to urobilinogen (**Figure 1A**). Of four strains that encode an annotated *bilR*, only *Mediterraneibacter gnavus* (formerly *Ruminococcus gnavus*) CC55_001C converted bilirubin to urobilinogen (**Figure 1B, top and Supplemental Figure 1A**). By contrast, the additional strains with an annotated *bilR* consumed bilirubin without concomitant production of urobilinogen despite picomolar detection sensitivity. To confirm this observation, we repeated the assay with the type strain of *M*. *gnavus* (ATCC 29149; hereafter *M. gnavus*) and with *Clostridium symbiosum* WAL-14163 (*C. symbiosum*), both of which encode BilR^23^. Consistent with the screen, *M. gnavus* reduced bilirubin to urobilinogen, whereas C*. symbiosum* consumed bilirubin but did not produce clearly detectable quantities of urobilinogen (**Figure 1C and 1D**).

To further evaluate if BilR from *M. gnavus* is sufficient for reduction of bilirubin to urobilinogen, we turned to heterologous expression in *Bacteroides thetaiotaomicron* VPI-5482 (*B*. *theta*). *B*. *theta* is a human gut bacterium that does not encode *bilR* and does not metabolize bilirubin (**Figure 1C and 1D**). *B*. *theta* with *M*. *gnavus bilR* stably integrated at the *attb2* site (*B*. *theta*^MgBilR+^) consumed bilirubin in culture but did not produce urobilinogen in clearly detectable quantities (**Figure 1C and 1D**). To extend this observation to a mammalian host, we colonized germ-free mice with a minimal, two-member bacterial community consisting of *B*. *theta*^WT^ and *E*. *coli* Nissle 1917, which deconjugates bilirubin glucuronide, or *B. theta*^MgBilR+^ and *E. coli* Nissle 1917. Mice colonized with *B. theta*^MgBilR+^ had lower levels of bilirubin in feces and cecal contents compared to mice colonized with *B*. *theta*^WT^, but urobilinogen was undetectable in both groups of mice (**Supplemental Figure 1B and 1C**). We detected urobilinogen in feces of specific pathogen-free mice using an identical procedure, demonstrating that our method is capable of detecting urobilinogen (**Supplemental Figure 1D**). These results establish that BilR is insufficient for the reduction of bilirubin to urobilinogen.

### Bilirubin reductase converts bilirubin to divinylurobilinogen

To identify the product of BilR, we inspected the LC–MS data from our screen for novel products. All strains that encode BilR produced a compound with m/z = 589.30, consistent with a 4-electron reduction of bilirubin (**Figure 1B, bottom**). We reasoned that this product could result either from the reduction of the vinyl groups in bilirubin (labeled blue in **Figure 1A**), producing mesobilirubin, or from the reduction of the methine bridges (labeled red in **Figure 1A**), producing divinylurobilinogen, a poorly characterized compound^9^. Comparison against an authentic mesobilirubin standard revealed that the product of these strains elutes at a different retention time than mesobilirubin (**Figure 1C and 1D, Supplemental Figure 1E**) and has a distinct MS/MS fragmentation pattern: Mesobilirubin generates one major fragment with m/z = 301.16, consistent with fragmentation at the central methylene bridge (**Figure 2A**, left), whereas the product of BilR-encoding strains generates an additional fragment with m/z = 466.23 (**Figure 2A**, right). This fragmentation pattern is consistent with the structure of divinylurobilinogen, in which the vinyl groups of the terminal pyrroles remain but the methine bridges have been reduced.

**Figure 2:**
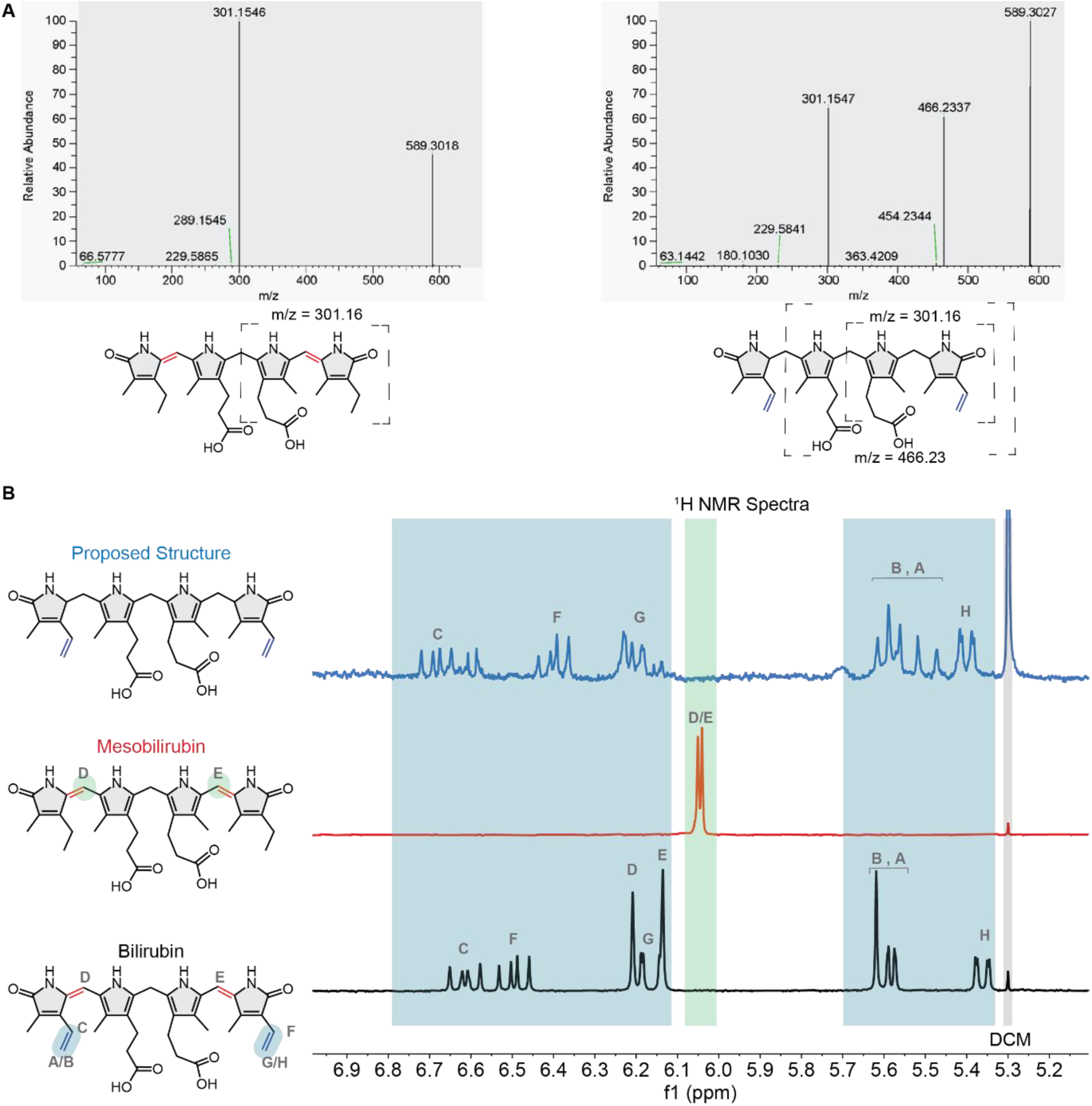
Bilirubin reductase produces divinylurobilinogen. **A.** Spectra from MS/MS fragmentation of NMR-verified mesobilirubin (left) or BilR product with m/z=589.30, generated by incubating *C. symbiosum* with 10 μM bilirubin for 24 hours (right). Fragmentation was performed with 10 eV for both compounds. **B.** ^1^H NMR of crude extract from *C. symbiosum* incubated with 10 μM bilirubin for 24 hours (top), mesobilirubin (middle), or bilirubin (bottom). Selected field of view displayed to highlight defining structures that differ across these three structures. Full spectra are included as supplemental figures 2, 3, and 4.

To further characterize the chemical structure of the product of BilR, we prepared crude extract of supernatant of *C. symbiosum* incubated with bilirubin and acquired an ^1^H NMR spectrum (**Figure 2B and Supplemental Figure 2**). We additionally acquired spectra of commercially obtained bilirubin and mesobilirubin for comparison (**Figure 2B and Supplemental Figure 3 and 4**). Although impurities in the crude extract affect the quality of the spectrum, the spectrum is consistent with the structure of divinylurobilinogen and not with those of mesobilirubin, bilirubin, or urobilinogen. Specifically, the *a*-vinyl protons of the terminal vinyl groups of bilirubin (labeled as C and F in **Figure 2B**) have been previously assigned as doublets of doublets, due to distinct couplings with the cis and trans β-vinyl protons, at observed values 6.6 ppm (dd, *J* = 17.3, 12.0 Hz) and 6.5 ppm (dd, *J* = 17.6, 11.5 Hz)^29^. This region of the spectrum of the extract also contains resonances corresponding to *a*-vinyl protons, with changes in chemical shift consistent with differences in the conjugation systems of the vinyl groups of bilirubin and divinylurobilinogen (6.6 ppm [dd, *J* = 17.7, 12.1 Hz] and 6.4 ppm [dd, *J* = 17.6, 11.6 Hz]). Resonances corresponding to the β-vinyl protons (labeled as A/B and G/H in **Figure 2B**) are also present in this region. Thus, the extracted compound contains terminal vinyl groups. Finally, the spectra of bilirubin and mesobilirubin contain resonances corresponding to methine protons (labeled as D and E in **Figure 2B**) as singlets at 6.13 ppm and 6.21 ppm for bilirubin and shifted upfield to 6.04 ppm and 6.05 ppm for mesobilirubin. The spectrum of the extracted compound does not contain resonances in either of these regions. We were unable to confirm the appearance of the methylene protons because the corresponding regions of the spectrum are difficult to interpret due to other components of the extract (**Supplemental Figure 2**). Nevertheless, these results are consistent with the absence of the methine bridges in the extracted compound. Together, our data establish that BilR only reduces the methine bridges in bilirubin to produce divinylurobilinogen, a compound whose existence had been previously proposed but never confirmed^9^.

**Figure 3:**
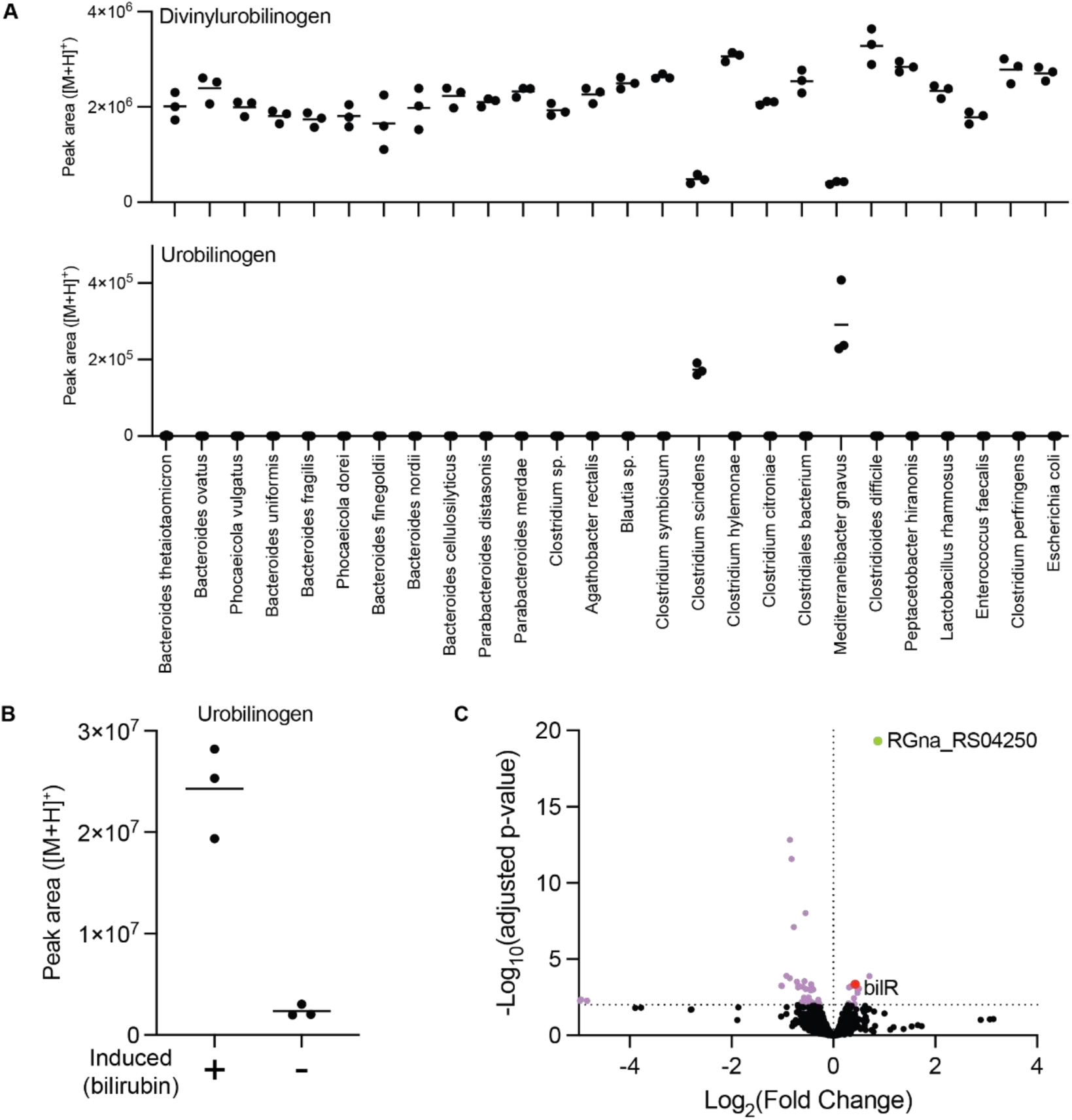
Identification of bilinoid vinyl reductase. **A.** Quantified peak area of remaining divinylurobilinogen (top) or urobilinogen (bottom) from indicated gut bacteria incubated with 10% spent media containing divinylurobilinogen. Each dot represents an individual bacterial culture. **B.** Substrate induction assay: quantified peak area of urobilinogen from *M. gnavus* pre-incubated with bilirubin (+) or DMSO (–) and then incubated with 10 μM bilirubin. Each dot represents an individual culture. **C.** Differential gene expression of *M. gnavus* incubated with 5 μM bilirubin for 30 minutes. Genes with statistically significant differences in expression (adjusted p-value < 0.01, as determined by DESeq2) are highlighted in purple. Data in A and B are represented as mean and individual values from replicate cultures.

**Figure 4:**
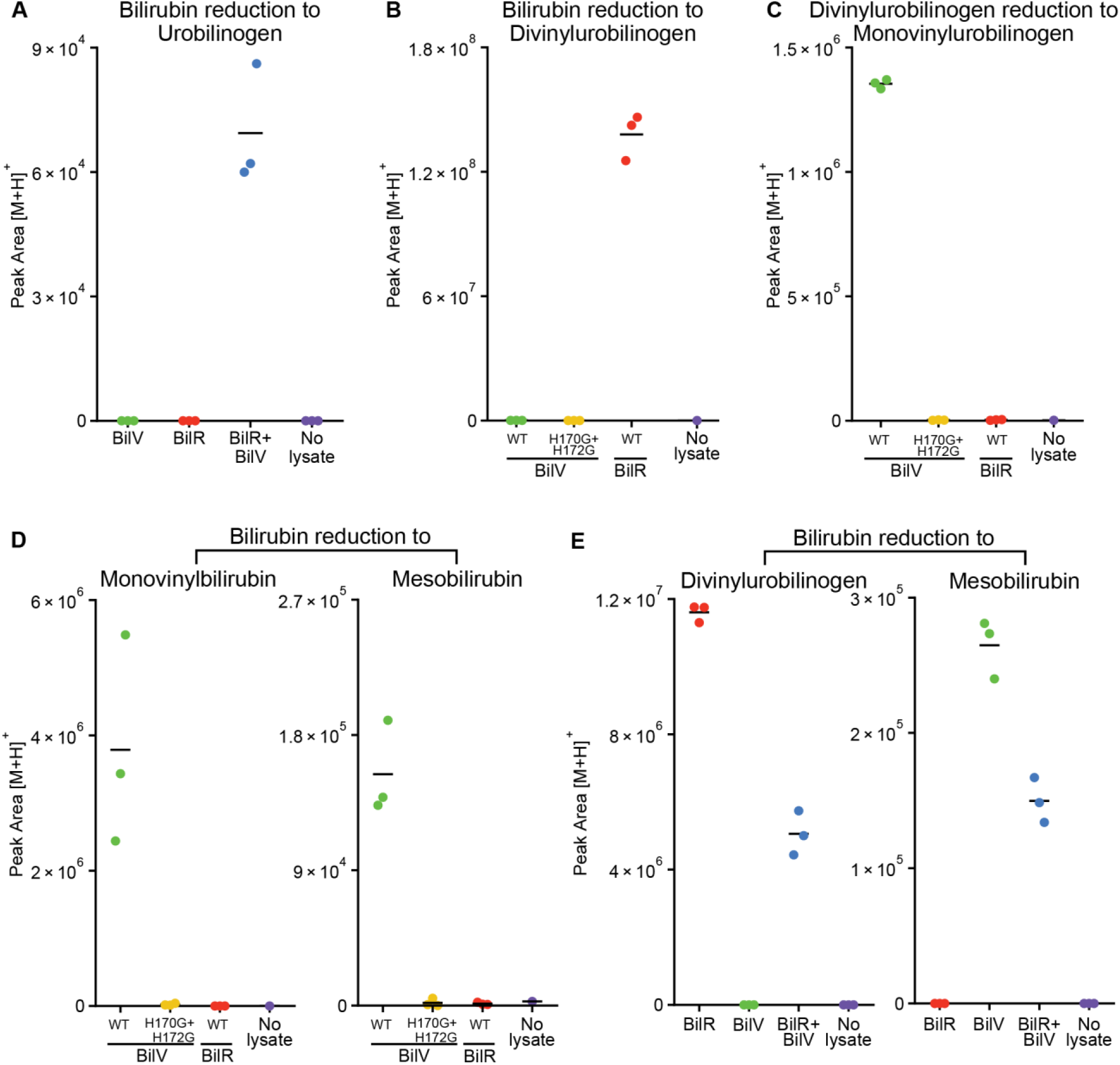
Bilinoid vinyl reductase reduces the vinyl groups in bilirubin and divinylurobilinogen. **A.** Quantified peak area of urobilinogen (m/z=593.33, retention time=2.6 min) from lysates of *E. coli* expressing BilR or BilV or from a mixture of the two lysates, each incubated with 500 nM bilirubin for 6 hours. **B.** Quantified peak area of divinylurobilinogen (m/z=589.30, retention time=2.5 min), from lysates of *E. coli* expressing the corresponding protein and incubated with 50 μM bilirubin for 6 hours. BilV H170G+H172G contains mutations of two histidine residues predicted to be located in the active site and take part in catalysis. **C.** Quantified peak area of c mpound with m/z=591.31 and retention time=2.5 min, consistent with monovinylurobilinogen, from lysates of *E. coli* expressing the corresponding protein and incubated with 5% spent media containing divinylurobilinogen for 6 hours. **D.** Quantified peak area of **(left)** compound with m/z=587.27 and retention time=2.5 min, consistent with monovinylbilirubin, and **(right)** mesobilirubin (m/z=589.30, retention time=6.7 min) from lysates of *E. coli* expressing the corresponding protein and incubated with 50 μM bilirubin for 6 hours. **E.** Quantified peak area of (**left**) divinylurobilinogen (m/z=589.30, retention time=2.5 min) or (**right**) mesobilirubin (m/z=589.30, retention time=6.7 min) from lysates of *E. coli* expressing BilR or BilV or from a mixture of the two lysates, each incubated with 500 nM bilirubin for 6 hours, except for the lysate mixtures, which were incubated with 50 µM bilirubin. All data are represented as mean and individual values of replicate lysates.

The product of BilR was previously evaluated using a fluorescence assay that relies on the strong fluorescence of urobilinogen and urobilin but not bilirubin when complexed with zinc^23^. (In such assays, urobilinogen is often first oxidized to the more stable urobilin.) We found that divinylurobilinogen, produced by incubation of *C*. *symbiosum* with bilirubin, also displays strong fluorescence in this assay (**Supplemental Figure 5A**), likely because divinylurobilinogen contains the same bilane core as urobilinogen and is therefore capable of complexing zinc.

To further evaluate the specificity of BilR for reduction of the methine bridges, we asked if *C. symbiosum* and *M*. *gnavus* can reduce mesobilirubin, in which the methine bridges of bilirubin are present but the vinyl groups are reduced to ethyl groups (**Supplemental Figure 5B**). Although *C. symbiosum* is unable to reduce bilirubin to urobilinogen, it can reduce mesobilirubin to urobilinogen (**Supplemental Figure 5B and 5C**). *M. gnavus* reduces both bilirubin and mesobilirubin to urobilinogen (**Supplemental Figure 5B and 5C**). These observations further confirm the specificity of BilR for reducing the methine bridges in bilirubin and mesobilirubin.

**Figure 5:**
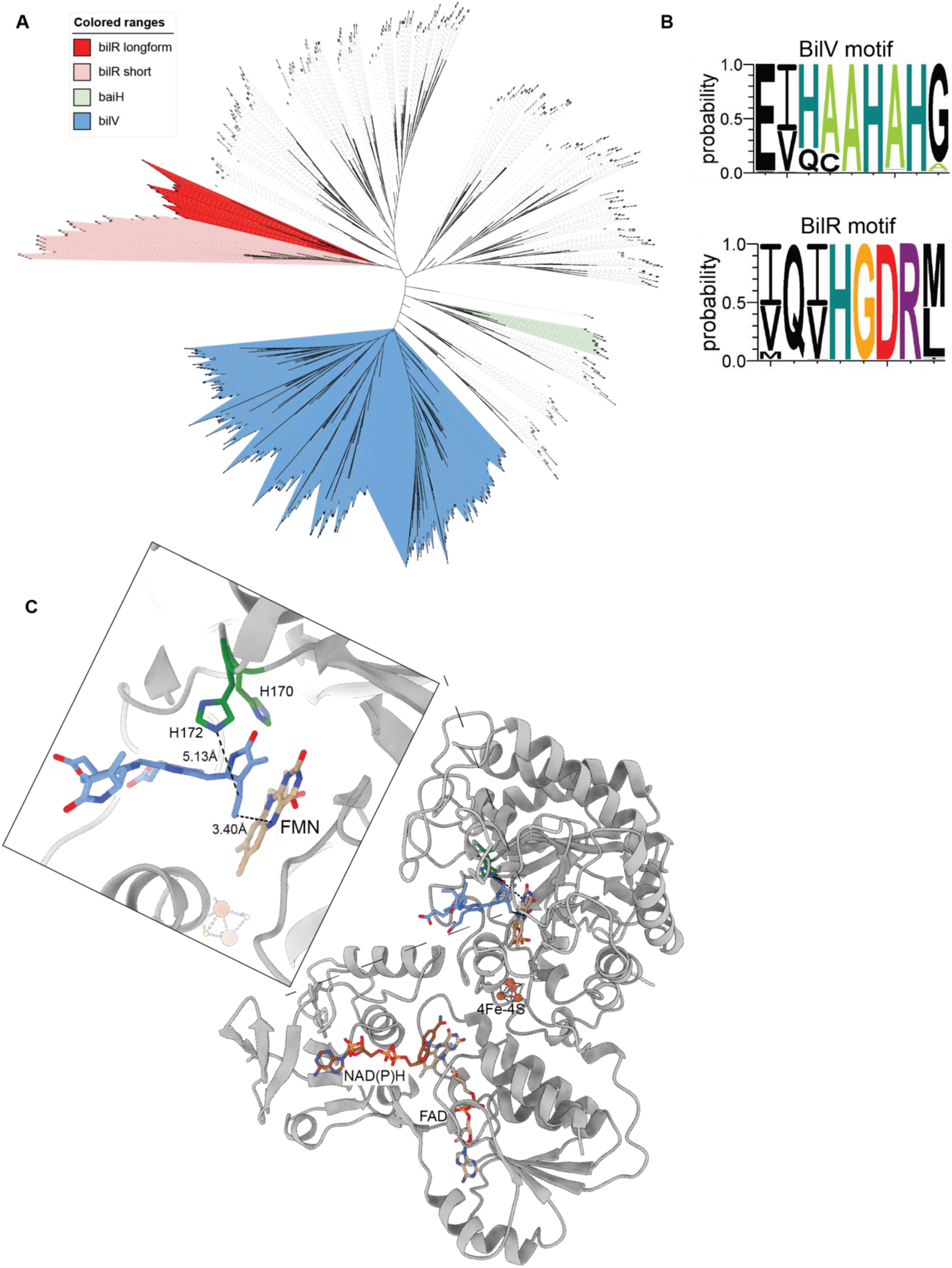
Delineation of the bilinoid vinyl reductase clade. **A.** Unrooted tree constructed from protein sequences of putati e bilinoid vinyl reductases (BilVs), bilirubin reductases (BilRs), and related enzymes from the Old Yellow Enzyme family. Putative BilVs are highlighted in blue, BilRs in pink and red, and 7-beta hydroxy-3-oxochol-24-oyl-CoA 4-desaturases (BaiHs) in green. **B.** Conserved sequence motifs in predicted binding pockets of putative BilVs (top) and BilRs (bottom). **C.** Structure of BilV from *M. gnavus* predicted by AlphaFold3 and modeled with the expected cofactors, FAD, FMN, and 4Fe-4S, and the substrates NAD(P)H and divinylurobilinogen. Inset highlights predicted binding pocket for divinylurobilinogen and distances between points of proton and hydride transfer required for the reduction reaction.

### Identification of a candidate bilinoid vinyl reductase

Our data suggested that at least one additional enzyme is required to reduce the vinyl groups in bilirubin. To identify bacteria capable of this reduction, we adapted our LC–MS screening approach. Specifically, because divinylurobilinogen is currently not commercially available, we incubated *C. symbiosum* with bilirubin to generate spent supernatant with divinylurobilinogen. We then supplemented the growth media of the 26 bacterial strains with 10% spent supernatant and monitored consumption of divinylurobilinogen and production of urobilinogen by LC–MS. Two strains*, Clostridium scindens* ATCC 35704 and *M. gnavus* CC55_001C, consumed divinylurobilinogen and produced urobilinogen (**Figure 3A**). These results confirm that *M*. *gnavus* is capable of producing urobilinogen and identify *C*. *scindens* as a novel gut microbe that participates in bilirubin metabolism.

To identify the enzyme responsible for this transformation, we turned to transcriptional profiling, motivated by previous findings that many reductases in gut bacteria are transcriptionally upregulated by their substrate^30–32^. In a substrate induction assay (Methods), pre-incubation of *M. gnavus* with bilirubin for 30 minutes resulted in increased capacity to convert bilirubin to urobilinogen (**Figure 3B**), suggesting that BilR and the unknown reductase are transcriptionally upregulated in the presence of bilirubin. We selected bilirubin as the inducer for these assays and subsequent RNA-seq to afford more precise control over substrate concentrations compared to divinylurobilinogen, which was not available in pure form at the time of our studies. Next, we performed RNA-seq of *M. gnavus* incubated with 5 µM bilirubin or vehicle for 30 minutes. Among the upregulated genes was *bilR*, as expected. The most significantly upregulated gene was an uncharacterized gene (RS04250) annotated to encode an FAD-dependent oxidoreductase with sequence similarity to BilR (**Figure 3C, Supplemental Table 2**). This annotation is consistent with the proposed reaction. *C. scindens* ATCC 35704, which had also converted divinylurobilinogen to urobilinogen (**Figure 3A**), encodes a protein with high identity to RS04250 (pairwise identity = 87.7%, **Supplemental Figure 6**), whereas the strains that had failed to convert divinylurobilinogen to urobilinogen in our screen do not (no protein with identity > 40%). These observations suggested that RS04250 is the enzyme that reduces the vinyl groups in divinylurobilinogen. We refer to RS04250 as bilinoid vinyl reductase (BilV), for reasons described in further detail below.

### Bilinoid vinyl reductase reduces the vinyl groups in bilirubin and divinylurobilinogen

To test if BilV and BilR together are sufficient for conversion of bilirubin to urobilinogen, we used a gain-of-function approach in *E*. *coli*. A mixture of lysates of *E*. *coli* expressing BilV and of *E*. *coli* expressing BilR converted bilirubin to urobilinogen (**Figure 4A and Supplemental Figure 7A**). Levels of produced urobilinogen were low but clearly distinguishable from contributions of isotopologues of more oxidized intermediates. By contrast, neither of the individual enzyme lysates produced urobilinogen in detectable quantities (**Figure 4A and Supplemental Figure 7A**). We next evaluated the individual enzymes for turnover of different substrates to establish substrate and product specificities. Lysates of *E. coli* expressing BilR converted bilirubin to divinylurobilinogen, as expected, and did not metabolize divinylurobilinogen further (**Figure 4B and 4C, Supplemental Figure 7B and 7C**). Lysates of *E*. *coli* expressing BilV did not convert bilirubin to divinylurobilinogen but converted divinylurobilinogen, predominantly into a compound with m/z = 591.32, consistent with a 2-electron reduction (**Figure 4B and 4C, Supplemental Figure 7B and 7C**). When analyzed by LC–MS, this compound eluted at a different time than urobilin (**Supplemental Figure 7C**), which has the same mass, and produced a different fragmentation pattern. Based on these properties, we propose that this compound is monovinylurobilinogen and is produced by the reduction of one vinyl group in divinylurobilinogen, perhaps because our assay system is not sufficiently optimized to promote further reduction to urobilinogen. Re-inspection of LC–MS traces obtained from native gut bacteria incubated with bilirubin or divinylurobilinogen also revealed the presence of this feature, consistent with monovinylurobilinogen being an intermediate of the pathway (**Supplemental Figure 7D**). Mutation of two histidine residues in the predicted active site of BilV (H170G+H172G), as described in further detail below, abolished turnover of divinylurobilinogen (**Figure 4C and Supplemental Figure 7C**). Although BilV is predicted to contain a 4Fe4S cluster based on similarity to 2,4-dienoyl-CoA reductase^33^, reduction of divinylurobilinogen occurred under both aerobic and anaerobic induction conditions, likely because the 4Fe4S cluster is shielded from solvent.

**Figure 6:**
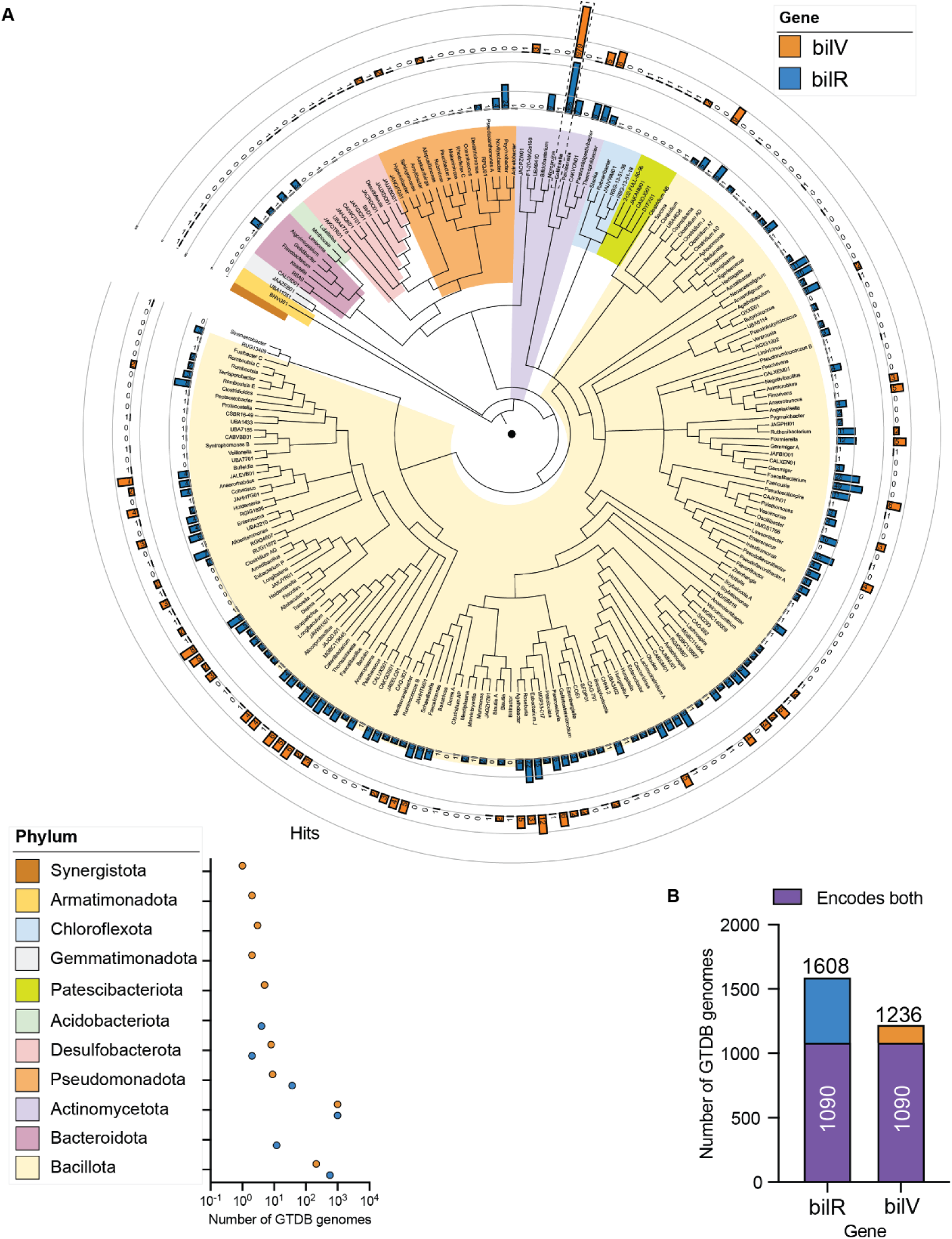
Taxonomic distribution of bilinoid vinyl reductase. **A.** Cladogram showing the relationships between taxa in the GTDB with detected bilirubin reductase (blue) or bilinoid vinyl reductase (orange), grouped by genus. Counts are plotted on a log scale. Bottom left: Number of genomes that encode bilinoid vinyl reductase (orange) or bilirubin reductase (blue), grouped by phylum. Box color represents phylum label for cladogram. **B:** Number of GTDB genomes that encode bilirubin reductase (blue), bilinoid vinyl reductase (orange), or both (purple).

**Figure 7:**
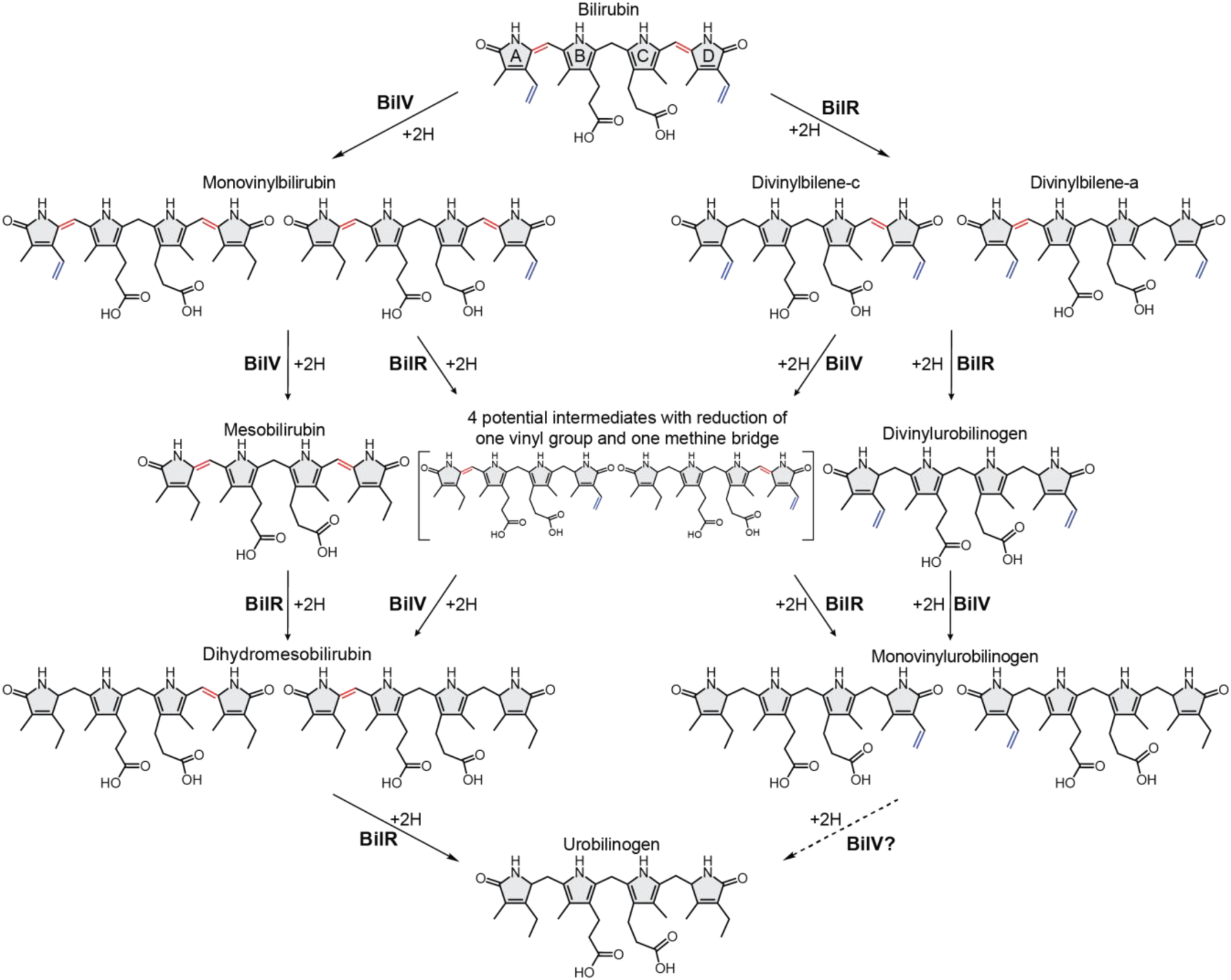
Pathway for metabolism of bilirubin to urobilinogen. Schematic includes reactions and intermediates detected in this study or previously or inferred from the double bond selectivities of BilR and BilV established in this study. Incubation of BilV with divinylurobilinogen results in formation of monovinylurobilinogen in lysate turnover assays (Figure 4C). The specificity of BilV for the vinyl groups, which are not equivalent, remains to be established. Both possible structures are shown for monovinylurobilinogen.

Bilirubin contains both vinyl groups and methine bridges and therefore could serve as a substrate for both BilR and BilV. Indeed, lysates of *E*. *coli* expressing BilV consumed bilirubin and produced a mixture of mesobilirubin, in which both vinyl groups are reduced, and monovinylbilirubin (also referred to as dihydrobilirubin), in which one of the vinyl groups is reduced (**Figure 4D, Supplemental Figure 7B and 8A**). Lysates of *E*. *coli* expressing BilR converted mesobilirubin to urobilinogen (**Supplemental Figure 8B and 8C**), consistent with the metabolism by strains that natively encode BilR (**Supplemental Figure 5**). These results suggested that the biochemical pathway for conversion of bilirubin to urobilinogen is branched. Indeed, mixing lysates of *E*. *coli* expressing BilV and of *E*. *coli* expressing BilR with bilirubin at higher concentrations (50 µM) led to appearance of both divinylurobilinogen and mesobilirubin (**Figure 4E and Supplemental Figure 8D**). The LC–MS traces also contained multiple additional, uncharacterized features with m/z = 589.30, which would be consistent with reduction of one vinyl group and one methine bridge, as well as with m/z = 587.29 and m/z = 591.32, which would be consistent with reduction of one of the double bonds and three of the double bonds, respectively (**Supplemental Figure 8E**). Thus, BilV performs the reduction of the vinyl groups in multiple compounds in the bilinoid family, including bilirubin and divinylurobilinogen. We therefore call this enzyme bilinoid vinyl reductase (BilV) and suggest that BilR should be renamed to bilinoid methine reductase. More broadly, our studies establish that BilV and BilR together are sufficient for conversion of bilirubin to urobilinogen and that the pathway for conversion of bilirubin to urobilinogen is branched.

### Delineation of the bilinoid vinyl reductase clade

BilV has homology to BilR and additional members of the Old Yellow Enzyme (OYE) family^34^. The sequence contains a predicted TIM barrel domain, a flavodoxin-like domain, and an NAD(P)H-binding domain similar to long-form BilR (previously also referred to as clade 1 BilR)^23^. In a phylogenetic tree of proteins homologous to BilV from *M. gnavus* (BilV^Mg^), both BilV^Mg^ and the putative BilV from *C. scindens* group in a single clade that is distinct from the clade formed by BilR and from other clades in the OYE family, such as the clade of BaiH enzymes that perform the desaturation step in the 7-dehydroxylation of bile acids^32^ (**Figure 5A**). Structures of BilR and BilV^Mg^ predicted by AlphaFold3 are highly similar to each other (**Supplemental Figure 9A**), including predicted binding sites for a flavin adenine dinucleotide (FAD) cofactor, which accepts a hydride from NAD(P)H, a flavin mononucleotide (FMN) cofactor, which performs the reduction of the substrate, and a 4Fe-4S cluster, which mediates electron transfer between the flavin cofactors. The overall structural similarity is consistent with a similar catalytic mechanism.

To identify determinants of substrate specificity for BilV and BilR, we further examined the sequences for differences in the active sites. All proteins in the BilV clade have a conserved AHAH motif (residues 169-172, *M*. *gnavus* numbering, **Figure 5B**) in a loop predicted by AlphaFold3 to be part of the substrate binding pocket adjacent to the FMN cofactor (**Figure 5C**). In BilR, the corresponding loop is three amino acids shorter and contains a conserved aspartate and arginine (D166 and R167, *M. gnavus* numbering, **Figure 5B, Supplemental Figure 9A and B**). These residues have been proposed to contribute to catalysis by donating the proton required to complete the reaction. In BilVs, these residues are replaced by two histidines from the conserved motif (H170 and H172, *M. gnavus* numbering). The positioning of these histidines together with the expansion of the corresponding loop changes the shape of the active site, which likely explains the difference in substrate specificity between BilV and BilR. In a structure of BilV bound to divinylurobilinogen predicted by AlphaFold3, the vinyl group of the A ring is positioned within hydrogen transfer distance of N5 of the FMN cofactor and the side chain of H172, consistent with transfer of a hydride and a proton during catalysis (**Figure 5C**). Mutation of H170 and H172 to glycines (BilV^Mg^ H170G+H172G) abolished conversion of divinylurobilinogen to monovinylurobilinogen and conversion of bilirubin to mesobilirubin in our *E*. *coli* lysate assay, establishing the importance of these histidines for the activity of BilVs (**Figure 4C and 4D**). Together, our results establish conserved sequence features to identify bilinoid vinyl reductases.

### Taxonomic distribution of bilinoid vinyl reductase

To define the taxonomic distribution of BilV, we generated a hidden Markov model from BilVs found in type strains of common human gut bacteria and queried predicted protein-coding amino acid sequences for 732,475 bacterial genomes in the Genome Taxonomy Database (GTDB, v226)^35^. Over 1,200 of these strains encode a putative BilV (**Figure 6A**). Similarly to BilR, most strains belong to the Bacillota and Actinomycetota. BilV is particularly prevalent in species of the genus Collinsella, with 979 of 1058 Collinsella genomes queried (92.5%) encoding BilV. BilV is also encoded across diverse members of the Bacillota but absent in the Bacteroidota.

Because our screens revealed that some bacteria carry out only one of the steps in the reduction of bilirubin to urobilinogen, we next evaluated co-occurrence of *bilV* and *bilR* in bacterial genomes. For this purpose, we used an analogous approach to identify genomes in the GTDB (v226) that encode BilR (**Figure 6A**). We found that ∼68% of strains that encode BilR also encode BilV (**Figure 6B**). Conversely, 88% of strains that encode BilV also encode BilR (**Figure 6B**). 930 of 1090 strains that encode both enzymes again belong to the genus Collinsella, and conversely 930 of 1058 Collinsella genomes queried (87.9%) encode both BilR and BilV. In many Collinsella genomes, BilR and BilV are encoded adjacent to each other in an arrangement consistent with expression as an operon. Thus, many Collinsella species are likely capable of transforming bilirubin to urobilinogen, implicating these species as dominant mediators of bilirubin metabolism. Outside of the Collinsella, some strains specialize in performing only one of the two reactions, as we observed for *C. symbiosum* and *C. scindens* in our screens. Most of the genomes that encode only BilR encode the short form of BilR, potentially suggesting functional diversification.

Finally, a query of 1,267 human metagenomes in the MetaQuery database revealed that BilV is both abundant (mean abundance = 0.022 gene copies per cell) and prevalent (97.5%) in the human gut microbiome (**Supplemental Figure 10A**)^36^. These results are consistent with observations that the conversion of bilirubin to urobilinogen is ubiquitous in the human population. Similarly, BilV is found in metagenomes of 81% of participants in the HMP2 cohort with a stringent workflow that only accounts for contributions of well-represented species (**Supplemental Figure 10B**). The taxonomic distribution of BilV in these metagenomes largely mirrors that in type strains (**Supplemental Figure 10C**). As in type strains, many species-level genome bins (SGBs) in these metagenomes encode both BilR and BilV, but some SGBs encode only BilR or only BilV, such that the major contributors to abundance of BilR and BilV in the gut microbiome are different: whereas the three major carriers of BilR are species of the genera *Faecalibacterium* and *Roseburia*, the major carriers of BilV are *M*. *gnavus* and species of the genera *Colinsella* and *Lachnospira* (**Supplemental Figure 10D-E**). The difference in BilV and BilR contribution is even more apparent within individual metagenomes. A single SGB accounts for over 90% of BilV abundance in 70% of BilV-positive samples, with a median of one contributing SGB, whereas BilR abundance is distributed across a median of seven SGBs per sample. These observations suggest that differences in microbiome composition across individuals can result in different carriages of BilR and BilV and therefore different abundances of products and intermediates of bacterial bilirubin metabolism.

## Discussion

Gut bacteria metabolize bilirubin in humans, but the transformations involved and the responsible bacterial species and enzymes have remained poorly characterized. Here, we demonstrate that conversion of bilirubin to urobilinogen requires distinct enzymatic steps, establish that BilR is selective for reduction of the methine bridges in bilinoids, and identify bilinoid vinyl reductase (BilV) as the enzyme responsible for reducing the vinyl groups (**Figure 7**). BilR and BilV are ubiquitous in healthy human microbiomes, supporting their assignment as prominent enzymes in bacterial bilirubin metabolism. While this manuscript was in preparation, an independent study also identified BilV via an orthogonal genomic approach^27^. Both our and this study provide complementary evidence that conversion of bilirubin to urobilinogen requires two enzymes, that BilV is responsible for reducing the vinyl groups in bilinoids, and that the pathway includes previously uncharacterized intermediates. Our study also includes the detection of multiple such intermediates and the conclusive annotation of divinylurobilinogen, whose existence had been first proposed in the 1970s^9^ and which likely had evaded detection because it is sensitive to oxygen and light and difficult to distinguish from other intermediates in the pathway with commonly used assays.

Because both BilR and BilV can accept bilirubin and intermediates of the pathway as substrates, the pathway for bilirubin metabolism is branched and proceeds via multiple distinct intermediates (**Figure 7**). These include mesobilirubin and divinylurobilinogen as well as candidate compounds with reduction of one or three of the four double bonds in bilirubin and with reduction of one methine bridge and one vinyl group. The concentrations of these intermediates and products and thus the output of bilirubin metabolism by a bacterial community then depend on the affinities of BilR and BilV for their substrates, the kinetics of the conversions, as well as the carriage of BilR and BilV in the community, as some species only encode one or the other enzyme, and the ability of species to take up compounds. For example, *C*. *scindens* encodes BilV but does not convert bilirubin to mesobilirubin.

BilV joins BilR, daidzein reductase, and BaiH, which performs the desaturation step in the 7-dehydroxylation of bile acids, as OYE family enzymes that catalyze reductive transformations in gut bacteria^32,37^. These reactions most likely serve to dispose of excess electrons from primary metabolism, akin to additional reductive transformations in gut bacteria^30,38^. Many such transformations are coupled to generation of ion gradients, either directly or via electron bifurcation, and thereby to ATP synthesis^30,38,39^. At present it is unclear if the mechanism of catalysis in OYEs, which involves electron transfer from NAD(P)H to the substrate, similarly allows for coupling to ATP synthesis. Because bilirubin is toxic to some Gram-positive species at elevated concentrations but not to Gram-negative species that have been tested^40,41^ and BilR and BilV are almost exclusively found in Gram-positive bacteria, it is also possible that BilR and BilV serve to detoxify bilirubin.

The assignment of BilV as the missing enzyme required for conversion of bilirubin to urobilinogen is supported by the production of urobilinogen upon mixing BilR and BilV with bilirubin in vitro as well as the ability of *C*. *scindens*, which encodes only *bilV* and not *bilR*, to convert divinylurobilinogen to urobilinogen. In our recombinant expression assay, however, BilV converts divinylurobilinogen primarily to monovinylurobilinogen. BilV does convert bilirubin to mesobilirubin, consistent with reduction of both vinyl groups. These observations likely owe to the asymmetric positioning of the vinyl groups. The vinyl group on ring A is conjugated to the carbonyl of the pyrrole and activated for reduction by a flavin-dependent reductase (**Figure 7**). By contrast, the vinyl group on ring D is unactivated in divinylurobilinogen and partially activated in bilirubin, through conjugation to ring C. BilV may be capable of reducing the ring D vinyl group in divinylurobilinogen with slower kinetics or under more optimized assay conditions, for example through a tautomer shift. It is formally possible that BilV is only capable of converting divinylurobilinogen to monovinylurobilinogen and that another enzyme converts monovinylurobilinogen to urobilinogen, although none of our studies suggest the existence of such an enzyme: no other putative oxidoreductases are found in genomic neighborhoods of *bilR* and *bilV*, even in organisms in which *bilR* and *bilV* are encoded adjacent to each other, and no putative oxidoreductases other than *bilR* and *bilV* were upregulated in *M*. *gnavus* upon addition of bilirubin. On the pathway level, conversion of bilirubin to urobilinogen may proceed more efficiently if reduction of the ring D vinyl group precedes reduction of the methine bridges, and monovinylurobilinogen may be produced in some quantities if reduction of the methine bridges occurs first. Indeed, monovinylurobilin was previously detected upon incubation of a human fecal isolate of *Clostridium perfringens* with bilirubin, likely after production of monovinylurobilinogen and subsequent non-enzymatic oxidation,^42^ and *C*. *scindens* and *M*. *gnavus* produce monovinylurobilinogen in addition to urobilinogen.

Many additional gaps remain in our understanding of bacterial bilirubin metabolism. The enzyme responsible for converting urobilinogen to stercobilinogen remains unknown, the stereospecificity of the transformations remains uncharacterized, and many additional candidate bilirubin metabolites that have been detected in metabolomics datasets^24^ remain to be chemically characterized. Much like the recent explosion in bacterial bile acids, we expect that many more bilirubin metabolites and transformations remain to be discovered, as do their roles in human health. Indeed, levels of bilirubin, urobilinogen, urobilin, other bilirubin derivatives, or BilR are associated with multiple inflammatory and metabolic conditions, including obesity, type 2 diabetes, and inflammatory bowel disease,^10,13–23,26,43–45^ as is abundance of the genus Collinsella, which contains many of the species that encode both BilR and BilV,^46–53^ and of members of the *Lachnospiraceae*, which have been recently implicated in bilirubin metabolism^25^. Our identification of species, enzymes, and metabolites involved in bacterial bilirubin metabolism now allows for precise mechanistic studies to define how this metabolism influences host physiology and contributes to disease.

## Methods

### Bacterial culturing

All strains used in this study are listed in Supplemental table 1. All bacterial culturing was performed in an anerobic chamber (Coy Laboratory Products) containing a gas mixture of 5% hydrogen and 10% carbon dioxide balanced with nitrogen unless otherwise stated. Strains were streaked from glycerol stocks onto brain heart infusion agar (BHI, Bacto) supplemented with 5 mg/L hemin, 2.5 µg/mL Vitamin K3, and 500 mg/L L-cysteine HCl (BHI+) unless otherwise stated and grown for 24-48 hours. Single colony isolates were then used to inoculate degassed BHI+ broth for experimentation. Plates were maintained for no longer than 10 days, stored under anaerobic conditions.

### Mice

All gnotobiotic mouse work was performed at the Gnotobiotic Core Facility at Harvard Medical School. All procedures were approved by the IACUC at Harvard Medical School under protocol IS00003227. Germ-free mice were bred and maintained in-house in climate-controlled rooms with a 12-hour light-dark cycle. Mice had access to food and water *ad libitum* and were housed in ISOcage P (Tecniplast) following weaning. Prior to the start the study, stool was collected from mice to confirm sterility. At 4-5 weeks of age, mice were gavaged with 100 μL of a 1:1 ratio of 10^8^ colony forming units of *E*. *coli* Nissle 1917 and either wild-type *Bacteroides thetaiotaomicron* or *Bacteroides thetaiotaomicron* expressing BilR (strain design described below). Stool was collected to confirm colonization on days 3, 7, and 14. Stool was collected for metabolomics on days 7 and 14. On day 14, mice were euthanized and cecal content was harvested for metabolomics analysis.

### Chemicals

The following chemicals were used for all assays: Bilirubin (Cayman Chemical, cat. 17161), Mesobilirubin (Santa Cruz Biotechnology, sc-263467), Urobilinogen (Medix Biochemica, 14684-37-8).

### Bacterial screen for reduction of bilirubin and derivatives

Thirty strains were analyzed (**Supplemental table 1**). Strains were recovered on YBHIS agar plates for 48 h at 37 C in a Coy anaerobic chamber (Coy Labs). YBHIS contained brain-heart-infusion powder (37 g/L), yeast extract (5 g/L), D-(+)-cellobiose (1 g/L), D-(+)-maltose monohydrate (1 g/L), cysteine (0.5 g/L), hemin (50 mg/L), and vitamin K1 (0.25 mg/L). All media were pre-reduced in the anaerobic chamber for at least 3 h before inoculation. Colonies were inoculated into 2 mL YBHIS in 96-deep-well plates, grown overnight, and then subcultured into 1 mL YBHIS for 3 h. Cultures were measured at OD600 using an Epoch2 spectrophotometer (BioTek Instruments) in clear, flat-bottom 96-well plates, and two parallel sets of 1 mL YCFA were then inoculated to obtain a final OD600 of 0.02 in each well. YCFA was prepared using 10 mL 10x salt solution plus acetate, 10 mL 10x casitone, 10 mL 10x yeast extract, 2 mL 5% cysteine, 1 mL 25% glucose, 36.5 μL 1 M MgSO4.7H2O, 8.2 μL 1 M CaCl2, 100 μL 0.5% hemin, 100 μL vitamin K1 (2.5 mg/mL), 10 μL ferrous sulfate, 1 mL ATCC vitamin mix, and autoclaved H2O to 100 mL final volume. Experimental YCFA cultures contained either 10 μM bilirubin (1% DMSO), 10% (v/v) spent medium containing divinylurobilinogen, or YCFA alone.

One plate set was processed to extract extracellular metabolites immediately, whereas the second was incubated for 24 h at 37 °C under anaerobic conditions before extraction. All experiments were performed in biological triplicate. For extracellular metabolite extraction, 225 μL of culture was centrifuged using an Eppendorf 5910 R centrifuge at 500 × g for 10 min at 4 °C. Then, 200 μL of cleared supernatant was transferred into 800 μL ice-cold methanol and centrifuged for 10 min. After centrifugation, 800 μL of supernatant was aspirated into a new plate. Plates were then sealed with aluminum tape and stored at -80°C until LC-MS analysis.

Bacterial strain identity was confirmed using 16S rRNA sequencing. Four strains from the library were excluded due to contamination, leaving twenty-six strains with reported values for these assays.

### Metabolite extraction

All metabolite extractions were performed under anaerobic conditions unless otherwise stated. All reagents and consumables where degassed at least 24 hours prior to use. All extractions occurred at room temperature. Cultures were vortexed prior to taking 1 mL and transferring it into a black microcentrifuge tube (Fisher Scientific, cat 50-403-867) to protect from light. Bacteria where pelleted at 16,000 × g for 10 minutes. Supernatant was transferred to a black microcentrifuge containing MS grade methanol (Thermo Scientific Chemicals, cat 047192.K2) to a final ratio of 1:4. Tubes were vortexed for 1 minute prior to centrifugation at 21,000 × g for 10 minutes. Supernatant was transferred to MS vials (MilliporeSigma, cat 29391-U) and tightly capped prior to removal from anaerobic chamber. Samples were then immediately run on LC-MS. For metabolite extraction from stool and cecal content of gnotobiotic mice, extraction occurred aerobically on ice. For stool, 80% methanol was added to 50 mg/mL. For cecal content 80% methanol was added to 25 mg/mL. Samples were homogenized in metal bead lysis tubes (MP Biomedicals, cat 6925050) in FastPrep®-24 Classic bead beating grinder and lysis system (MP biomedicals) for two rounds of 3.0 m/s for 30 seconds each. Samples were rested on ice for 2 minutes between homogenization rounds. Samples were centrifugation at 21,000 × g for 10 minutes. Supernatant was transferred to MS vials (MilliporeSigma, cat 29391-U) and immediately run on LC-MS.

### Liquid chromatography–Mass Spectrometry

Target compounds were detected by liquid chromatography–mass spectrometry using a Thermo Vanquish – Exploris 240 LCMS system. All data and instrument method files for this study are available at the Metabolomics Workbench^54^ under project PR003449. Briefly, samples were injected and separated on a Waters HSS T3 C18 2.1 x 100 mm column using 0.1% formic acid (A) and acetonitrile with 0.1% formic acid (B) as liquid phases and a gradient from 40% to 98% B over 2 minutes followed by a 6 minute 98% B hold and re-equilibration steps at 0.3 mL/min and 40 °C. Source parameters were as follows: positive ion: 3500 V, negative ion: 3000 V, sheath gas 50 (Arbitrary Units), aux gas 10 (Arbitrary Units), sweep gas 1 (Arbitrary Units), ion transfer tube temp 325 °C, vaporizer temp 350 °C. Scan properties were as followed: for both positive and negative ion mode, scan range 250 - 750 m/z, resolution 60,000, RF Lens 70%. Tandem MS/MS fragmentation was performed at energies of 10 eV, 20 eV, and 30 eV in a data-dependent manner for [M+H]^+^ consistent with mesobilirubin and divinylurobilinogen (589.3021 m/z with 1 m/z resolution), Orbitrap resolution 30,000, and normalized collision energy type. The presented fragmentation was performed with 10 eV because it allowed for the retention of the parent mass while visualizing the daughter fragments of interest. The mass spectrometer was cleaned and calibrated weekly using Pierce FlexMix calibration solution following instrument vendor guidelines. Urobilinogen was quantified after correcting for abundances of isotopologues of lower molecular weight intermediates with similar retention times (monovinylurobilinogen and divinylurobilinogen) using expected natural isotope abundances.

### Generation of *B*. *theta*^MgbilR+^

*B*. *theta*^MgbilR+^ was generated using previously described counter-selectable allelic exchange^55^. Briefly, 1 kb fragments upstream and downstream of one of the *attb2* sites from *Bacteroides thetaiotaomicron* VPI-5482 were amplified using the following primers: upstream homology arm: Forward 5ʹ-ACCGTAGTATTCGCCCAT; Reverse: 5ʹ-GGATTCGAACCCCGGG. Downstream homology arm: Forward: 5ʹ-CCTGTCTCTCCGCTGGAAA; Reverse: 5ʹ-ATATCCCGTACCTGACTTAACCG. Long-form bilR was amplified from *M*. *gnavus* ATCC 29149 using the following primers: Forward: 5ʹ-ATGAGATTATTAGAACCAATTAAAGTGGGA; Reverse: 5ʹ-TAAAACGTTTGCTGCCAGAT. Promoter pBfP1E4 and ribosomal binding site RBS4 were added to long-form bilR using overlap extension PCR^56^. The genome homology arms and MgbilR were assembled into pLGB13 using Gibson Assembly. The vector was transformed into *E*. *coli* S17λpir and then conjugated into *B*. *thetaiotaomicron*. Single-crossover integrants were selected on BHI+ agar plates containing 200 μg mL^−1^ gentamicin and 25 μg mL^−1^ erythromycin. Integrants were grown overnight in non-selective BHI+ broth and then plated on BHI+ agar plates containing 100 ng mL^−1^ anhydrotetracycline for counterselection. Candidates were screened for successful insertion using PCR, confirmed for sensitivity to erythromycin, and sequenced to confirm no mutations occurred during the cloning process.

### NMR Instrumentation

Nuclear magnetic resonance spectra were recorded on a 400 MHz spectrometer equipped with a 5mm AutoX One Probe. Chemical shifts are reported relative to residual solvent peaks in parts per million (CHCl3: ^1^H, 7.26). Acquired with 128 scans.

### Nuclear Magnetic Resonance of divinylurobilinogen

An overnight culture of *C*. *symbiosum* was diluted into 1 L of degassed BHI containing 10 µM of bilirubin protected from light. The culture was incubated anaerobically at 37 °C for 24 hours. After removal from the anaerobic chamber, the culture was spun down at 4300 × g for 10 minutes and supernatant was collected. In two 500 mL batches, the supernatant was decanted into a separatory funnel, and this aqueous layer was extracted with dichloromethane three times (3 x 50 mL). The combined organic phase was dried under reduced pressure. The crude material was resuspended in 40% acetonitrile in water (1 mL) and loaded onto an equilibrated Bakerbond™ octadecyl (C18) reverse phase pre-packed column (100 mg solid phase) (Cat#: Avantor JT-7020-01). The loaded column was washed three times with 40% acetonitrile in water (3 x 1mL) and then the product was eluted in 100% acetonitrile. Solvent was removed under reduced pressure. Resultant residue was taken up in deuterated chloroform for NMR analysis.

<u>Bilirubin Standard:</u> ^1^H NMR (400 MHz, CDCl3) δ 13.69 (s, 2H), 10.75 (s, 2H), 9.28 (s, 2H), 6.61 (dd, *J* = 17.3, 12.0 Hz, 1H), 6.50 (dd, *J* = 17.6, 11.5 Hz, 1H), 6.17 (m, 3H), 5.62 (ddd, *J* = 17.4,11.7, 1.2 Hz, 1H), 5.58 (dd, *J* = 10.5, 1.2 Hz, 1H), 5.36 (dd, *J* = 11.4, 2.2 Hz, 1H), 4.08 (s, 2H), 3.10 – 2.74 (m, 6H), 2.58 (d, *J* = 14.2 Hz, 2H), 2.20 – 2.10 (m, 9H), 1.99 (s, 3H).

<u>Mesobilirubin Standard:</u> ^1^H NMR (400 MHz, CDCl3) δ 13.65 (s, 2H), 10.59 (s, 2H), 9.15 (s, 2H), 6.05 (s, 2H), 4.07 (s, 2H), 3.06 – 2.73 (m, 6H), 2.61 – 2.53 (m, 2H), 2.48 (q, *J* = 7.6 Hz, 2H), 2.32 (q, *J* = 7.6 Hz, 2H), 2.16 (d, *J* = 0.8 Hz, 6H), 2.07 (s, 3H), 1.86 (s, 3H), 1.09 (dt, *J* = 22.5, 7.6 Hz, 6H).

<u>Crude extract annotated region:</u> ^1^H NMR (400 MHz, CDCl3) δ 6.65 (dd, *J* = 17.7, 12.1 Hz, 1H), 6.40 (dd, *J* = 17.6, 11.6 Hz, 1H), 6.21 (m, 1H), 5.60 (m, 1H), 5.52 (dd, 17.7, 0.6 Hz 1H), 5.40 (dd, *J* = 11.4, 2.0 Hz, 1H).

### Fluorescence assay for detection of bilirubin derivatives

This protocol was adapted from previous work^23^. Briefly, overnight cultures were diluted 1:100 into 50 mL of BHI+ containing 5 μM bilirubin protected from light. Cultures were incubated anaerobically at 37 °C for 24 hours. Cultures were then pelleted by centrifugation at 7000 x g for 10 minutes. Supernatant was collected and removed from the anaerobic chamber. Supernatants were then filtered with 0.22 μM filter into sterile glass bottles containing 1.25 mL of chloroform and protected from light. Bottles were inverted several times to aid the extraction. Using a glass Pasteur pipette, approximately 1 mL of chloroform was collected and transferred to a black microcentrifuge tube (Fisher Scientific, cat 50-403-867) to protect from light. Chloroform was then evaporated using an Eppendorf Vacufuge plus at room temperature. After evaporation, 600 μL miliQ water was added and vortexed for 1 minute. 400 μL was transferred to a fresh tube, to which 10 μL of 10% povidone-iodine was added, followed by 10 μL of 100 mM cysteine with vortexing in between. Finally, 400 μL of 545 mM zinc acetate dissolved in methanol was added and samples were vortexed for 1 minute. 100 μL from each sample was transferred in triplicate to a black 96-well plate and fluorescence intensity was recorded at 495 nm wavelength excitation and 525 nm wavelength emission (EnSight Multimode Plate Reader, PerkinElmer).

### Induction test

Testing for transcriptional upregulation of the candidate BilV was performed as outlined in Bae et al., 2025^31^. Briefly, three individual colonies of *M*. *gnavus* were inoculated into BHI broth for three independent replicates. Overnight cultures were diluted to OD 0.01 and allowed to grow into early exponential phase (OD 0.1-0.2). Cultures were then split, and received either DMSO (uninduced) or bilirubin (dissolved in DMSO) at a final concentration of 5 µM (induced). After 30 minutes of incubation, samples were spun down and washed 3 times with PBS. Finally, samples were suspended in 1 mL of PBS, split into two tubes, and treated with DMSO or 5 µM of bilirubin. After 30 minutes of incubation, samples were processed as outlined in metabolite extraction section and divinylurobilinogen levels were measured by LC–MS.

### RNA-seq

Three individual colonies of *M*. *gnavus* were inoculated into BHI broth, for three independent replicates. Overnight cultures were diluted to OD 0.01 and allowed to grow into early exponential phase (OD 0.1-0.2). Cultures were then split and received either DMSO or bilirubin (dissolved in DMSO) at a final concentration of 5 µM. After 30 minutes of incubation, samples were processed as outlined in Protocol 5 of the Qiagen RNAprotect Bacteria Reagent handbook. Briefly, 2x volume of RNAprotect Bacteria Reagent was added to each culture. Samples were pelleted and underwent enzymatic lysis, proteinase K digestion, and mechanical disruption using a Retch MM400 bead beater. Samples were DNase treated with DNase (RNase free, Invitrogen). RNA sequencing library preparation using rRNA depletion and Illumina sequencing was performed by SeqCenter (Pittsburgh, PA). Library preparation was performed using the Stranded Total RNA Prep Ligation with Ribo-Zero Plus kit (Illumina) and 10bp unique dual indices (UDI). Libraries were sequenced on a NovaSeq X Plus (Illumina), producing paired end 150bp reads. Demultiplexing, quality control, and adapter trimming was performed with bcl-convert (v4.2.4). Reads were aligned to the *M*. *gnavus* ATCC 29149 genome (RefSeq ID: GCF_009831375.1) using Bowtie2 (version 2.5)^57^ with default parameters and aligned features were counted using featurecounts (version 2.24)^58^. Differential expression analysis was performed using DESeq2 (version 1.50)^59^ with default parameters.

### Generation of *E*. *coli*^BilV^, *E*. *coli*^BilV_H170G+H172G^, *E*. *coli*^BilR^ and turnover assays

BilV was amplified from *M*. *gnavus* ATCC 29149 using the following primers: Forward: 5ʹ-ATGAATACATATCCAAAACTCTTC; Reverse: 5ʹ-TTATAAATTCAATGCAGTCTG. BilR was amplified as described above. The amplified genes were inserted into pET28a using Gibson Assembly and transformed into MAX Efficiency Stbl2 Competent Cells (Invitrogen, 10268019). A construct encoding H170G/H172G BilV was generated by site directed mutagenesis of the BilV construct using the following primers: Forward: 5ʹ-CAGGCGGACTCCTTGGCGGATTC; Reverse: 5ʹ-CGCCAGCGGCGTGAATTTCC. Constructs were confirmed using Nanopore sequencing (Quintara). The resulting constructs where then transformed into Rosetta(DE3) pLysS competent cells (Millipore Sigma, 70956-M). For each replicate, 5 colonies were selected and inoculated into LB for overnight growth for a total of three replicates. Overnight cultures where then diluted 1:100 into 15 mL of LB and allowed to grow into late exponential phase (OD 0.6-0.8). Protein expression was induced with 100 μM IPTG and cultures were incubated for 16 hours at 25°C. Cultures were then pelleted and media was removed. Pellets underwent three cycles of freeze-thaw on dry ice with ethanol baths to lyse bacterial cells. Lysates were moved into the anaerobic chamber, resuspended in 5 mL of degassed BHI, and diluted 1:10 (Figure 4B-D, Supplemental 7B-C, and Supplemental 8A) or 1:50 (Figure 4A, 4E, Supplemental 7A, Supplemental 8B-E). For experiments with individual lysates of *E*. *coli* expressing BilR or of *E*. *coli* expressing BilV, 500 µL of diluted lysate was incubated with 5% spent media containing divinylurobilinogen or 500 nM – 50 μM bilirubin in the presence of 5mM NADH. For experiments with mixtures of lysates of *E*. *coli* expressing BilR and of *E*. *coli* expressing BilV, 250 µL each of the diluted lysates were mixed and incubated with 5% spent media containing divinylurobilinogen or 500 nM – 50 μM bilirubin in the presence of 5mM NADH.

### Bioinformatic analyses

AlphaFold3 was used to model the structures of BilR and BilV from *M*. *gnavus* with flavin mononucleotide and to model a structure of BilV with flavin mononucleotide, flavin adenine dinucleotide, nicotinamide adenine dinucleotide phosphate, 4Fe-4S, and divinylurobilinogen. Models were visualized in ChimeraX-1.11.1.

Construction of the phylogenetic tree of OYEs (figure 4A): RS04250 was used to search the Reference Protein database using BLASTp, using the default parameters except for an increase in results to 1000 sequences (minimum E-value = 1xe^-100^). Separate searches were performed for a representative long form (Uniprot ID: A0A829NF98) and short form (Uniprot ID : E7GT89) BilR. For each BilR form, the top 250 hits were selected and added to the list resulting in 1500 sequences. The 1500 sequences were collapsed to 1490 non-redundant 1490 sequences, which were then aligned using Clustal Omega using the default parameters. The resulting alignment was used to construct a tree using Geneious Tree Building Neighbor-joining method with bootstrap resampling with 100 replicates. The resulting tree was visualized using interactive tree of life (iTOL).

Evaluation of phylogenetic distribution of BilV and BilR (Figure 5A): RS04250 was used to search a custom database of genomes containing representative type strains found in the microbiome with deposited genomes on Integrated Microbial Genomes and Microbiomes hosted by the Joint Genome Institute^60^. High confidence hits (E-value = 0.0) were then aligned using a multisequence alignment (Clustal Omega)^61^. The resulting alignment was then used to build a Hidden Markov Model using HMMER (version v3.4) hmmbuild using the default parameters. For BilR, two hidden Markov models were constructed using the sequences provided in Supplemental Table 1 of Hall et al., 2024 for short form BilR and long form BilR using hmmbuild using the default parameters.

The constructed model was used to query the Genome Taxonomy Database (GTDB) Release 10-RS226 amino acid sequences using hmmsearch with an E-value threshold of 1e^-5^, and otherwise default parameters. The results were then filtered for an E-value of less than 1e^-100^ and for the presence of the conserved motif ‘AHAH’ for divinylurobilinogen or ‘HGDR’ for BilR. Resulting hits were then mapped back to their corresponding genomes and collapsed to the genus level. These hit counts were plotted onto a condensed version of the phylogenetic tree from GTDB, with only genera that contained either bilR or bilV kept for visualization. This tree and the hit counts were then visualized using interactive tree of life (iTOL).

Evaluation of prevalence and abundance of bilV in human metagenomes: RS04250 was used to search the Metaquery^36^ database using the following search parameters: min percent identity= 60%, maximum evalue: 1e-40, min query alignment coverage = 70% (default), min target alignment coverage = 70% (default). The resulting hit table was inspected to confirm that all hits contained the conserved ‘AHAH’ motif of BilV. The analyzed set of shotgun metagenomes (MGX) from HMP2^62^ consisted of 1638 stool metagenomes from 130 participants sampled longitudinally. Taxonomic profiles were generated with MetaPhlAn v4.0.6^63^ against the mpa_vOct22_CHOCOPhlAnSGB_202403 marker database, and functional profiles were generated with HUMAnN v4.0.alpha.1^64^. Reference sequences were the characterized enzymes from this study: BilV of *Mediterraneibacter gnavus* and *Clostridium scindens*, and BilR of *M. gnavus*, *Clostridium symbiosum*, and *Clostridioides difficile*^23^. These protein sequences were searched against the ChocoPhlAn catalog (build v202210_202403) with tBLASTn (BLAST+ 2.17.0; evalue 1e-3, max_target_seqs 5000). Cutoffs of amino acid identity ≥ 50%, query coverage ≥ 80%, and subject coverage ≥ 80% resulted in 943 unique gene IDs that collapsed to 417 unique HUMAnN features. Within the HMP2 cohort, there were 94 HUMAnN observed gene families (21 *bilV* and 73 *bilR*). Examining the HUMAnN features, 100% of the HMP2 *bilV* total abundance and 100% of the *bilR* total abundance are contributed by gene families containing the diagnostic active site motif (AHAH for BilV and HGDR for BilR). Stratified abundances for the assigned HUMAnN features were summed per species-level genome bin (SGB) to give the contribution per SGB of *bilV* and *bilR*. In the phylogenetic tree, SGBs were mapped to GTDB r226 lineages.

## Acknowledgements

We thank Michael Fischbach, Jakub Rajniak, Elizabeth Hart, and all members of the Jost and Rakoff-Nahoum labs for helpful discussions, the Analytical Chemistry Core at Harvard Medical School for providing LC-MS instrument access and expertise, and Kathryn Durben and the HMS Gnotobiotic Core Facility for gnotobiotic mouse husbandry. This work was supported in part by the National Institutes of Health (grants R00GM130964 and DP2GM154152 to MJ, R01AI148752 to SW, T32DK135449 to EH, DP2GM136652, R01DK138023, R01AI158814, R01AI126915 and R01AI171100 to SR-N, R24DK110499 and P30DK043351 to CH), a Helen Hay Whitney Foundation fellowship (to VMM), the Kenneth Rainin Foundation (to MJ), the Resnek Center for PSC Research (to EH and SRN), a Career Award for Medical Scientists from the Burroughs Wellcome Fund (to SR-N), a Pew Biomedical Scholarship (to SR-N), and a Graduate Research Fellowship from the National Science Foundation (DGE 2630152 to DHS). Availability of the data at the NIH Common Fund’s National Metabolomics Data Repository is supported by NIH grants U2C-DK119886 and OT2-OD030544.

## Author Contributions

BJR performed LC-MS-based characterization of bacterial bilirubin metabolism, isolation and characterization of divinylurobilinogen, identification and characterization of BilV, bioinformatic analyses, analyzed experimental data, and drafted the manuscript together with MJ. EJH performed the bacterial library screens and contributed to LC-MS-based characterization of metabolism assisted by JL. VMM performed and analyzed the NMR with supervision from SW. JMC and MG constructed the *E*. *coli* gain-of-function strains and assisted with bacterial culturing. NHD, KNT, and EJH performed mapping of BilR and BilV in HMP2 data with supervision from CH. DHS contributed to characterization of activities of BilR and BilV in recombinant expression assays. MJJ developed and oversaw all LC-MS. SRN and MJ supervised all work. BJR, EJH, SRN, and MJ conceived the project. BJR, EJH, DHS, SRN, and MJ revised the manuscript. All authors provided feedback on the manuscript.

## Data Availability

Sequencing reads from RNA-seq of *M*. *gnavus* with or without bilirubin are available at the NCBI Sequence Read Archive (SRA) under BioProject accession number PRJNA1477165. Processed counts are included as Supplementary Table 3. LC–MS, LC–MS/MS, and NMR data are available at the NIH Common Fund’s National Metabolomics Data Repository (NMDR) website, the Metabolomics Workbench (https://www.metabolomicsworkbench.org), under Project Number PR003449 (https://dx.doi.org/10.21228/M81R6N098; study IDs ST005336, ST005337, ST005338, and ST005339).

---

**Supplemental Figure 1:**
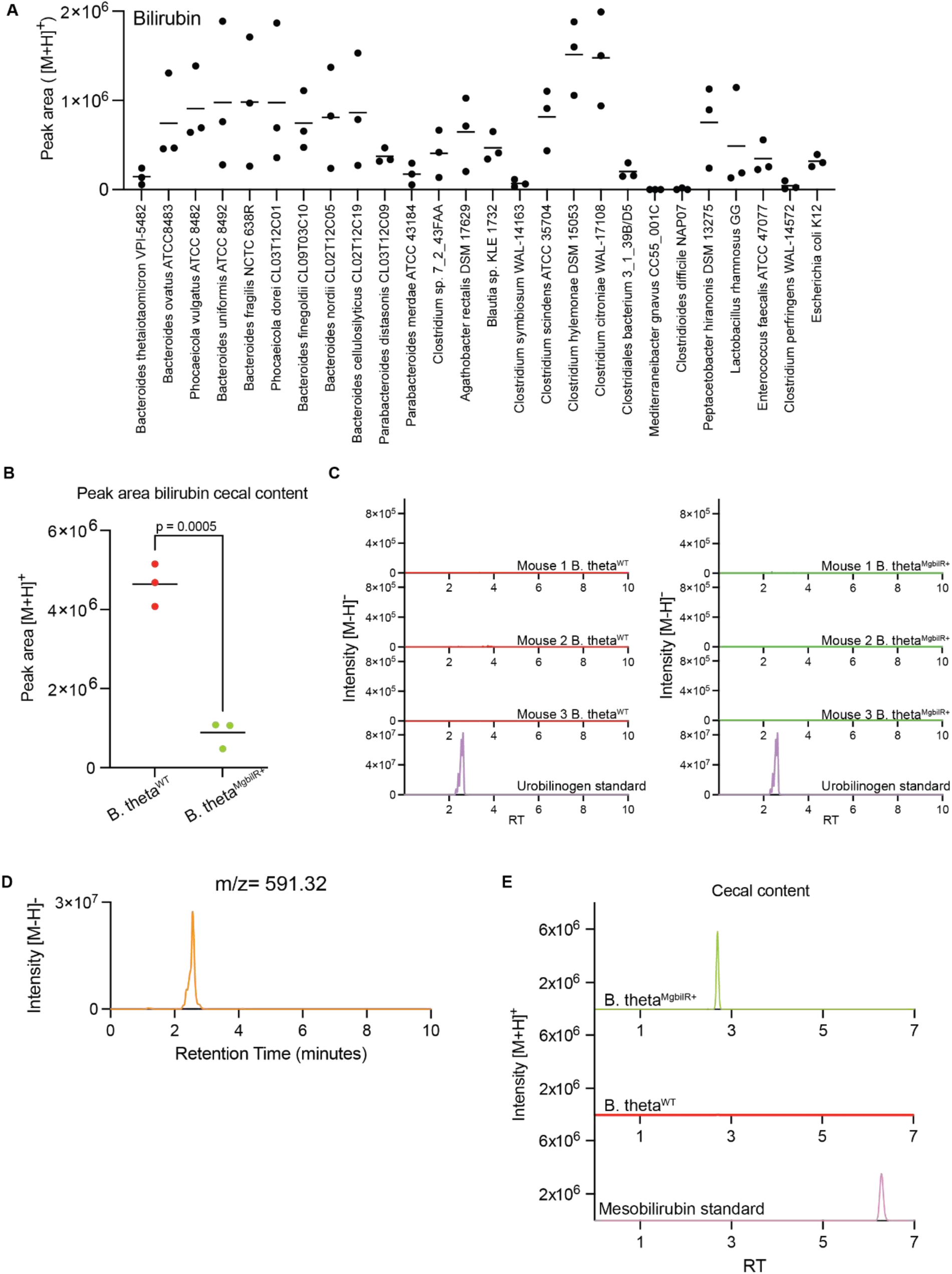
BilR is insufficient to convert bilirubin to urobilinogen. **A.** Quantified peak area of remaining bilirubin (standard-verified) from indicated gut bacteria incubated with 10 μM bilirubin for 24 hours. Each dot represents an individual bacterial culture. **B.** Quantified peak area of standard verified bilirubin in mice colonized with *E*. *coli* Nissle 1917 and either *B*. *theta*^WT^ (red) or *B*. *theta*^MgbilR+^ (green) for 14 days. *p*-value derived from unpaired t-test. **C.** Extracted ion chromatograms for urobilinogen in feces of mice colonized with indicated *B*. *theta* strains. Each plot represents feces from an individual mouse. **D.** Extracted ion chromatogram for urobilinogen in feces of specific pathogen-free mouse. **E.** Representative extracted ion chromatograms for m/z=589.30 in positive ion mode in cecal contents of mice colonized with *E*. *coli* Nissle 1917 and either *B*. *theta*^WT^ (red) or *B*. *theta*^MgbilR+^ (green) for 14 days. Each plot represents cecal content of an individual mouse. Bottom plot is a commercially obtained mesobilirubin standard. Data in A and B are represented as mean and individual values from replicate cultures (A) or individual animals (B).

**Supplemental Figure 2:**
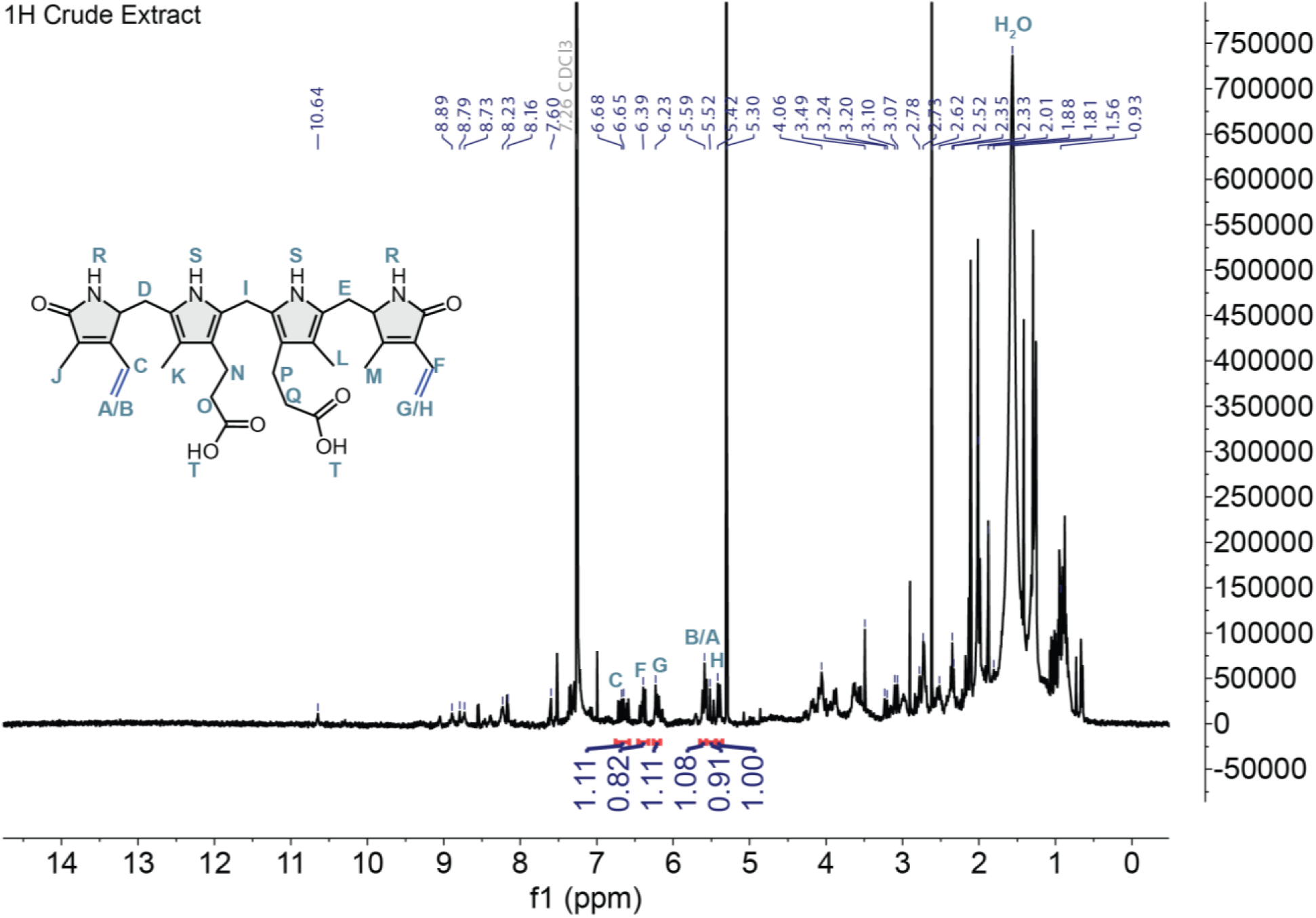
Full ^1^H NMR spectrum of crude extract from *C. symbiosum* incubated with bilirubin.

**Supplemental Figure 3:**
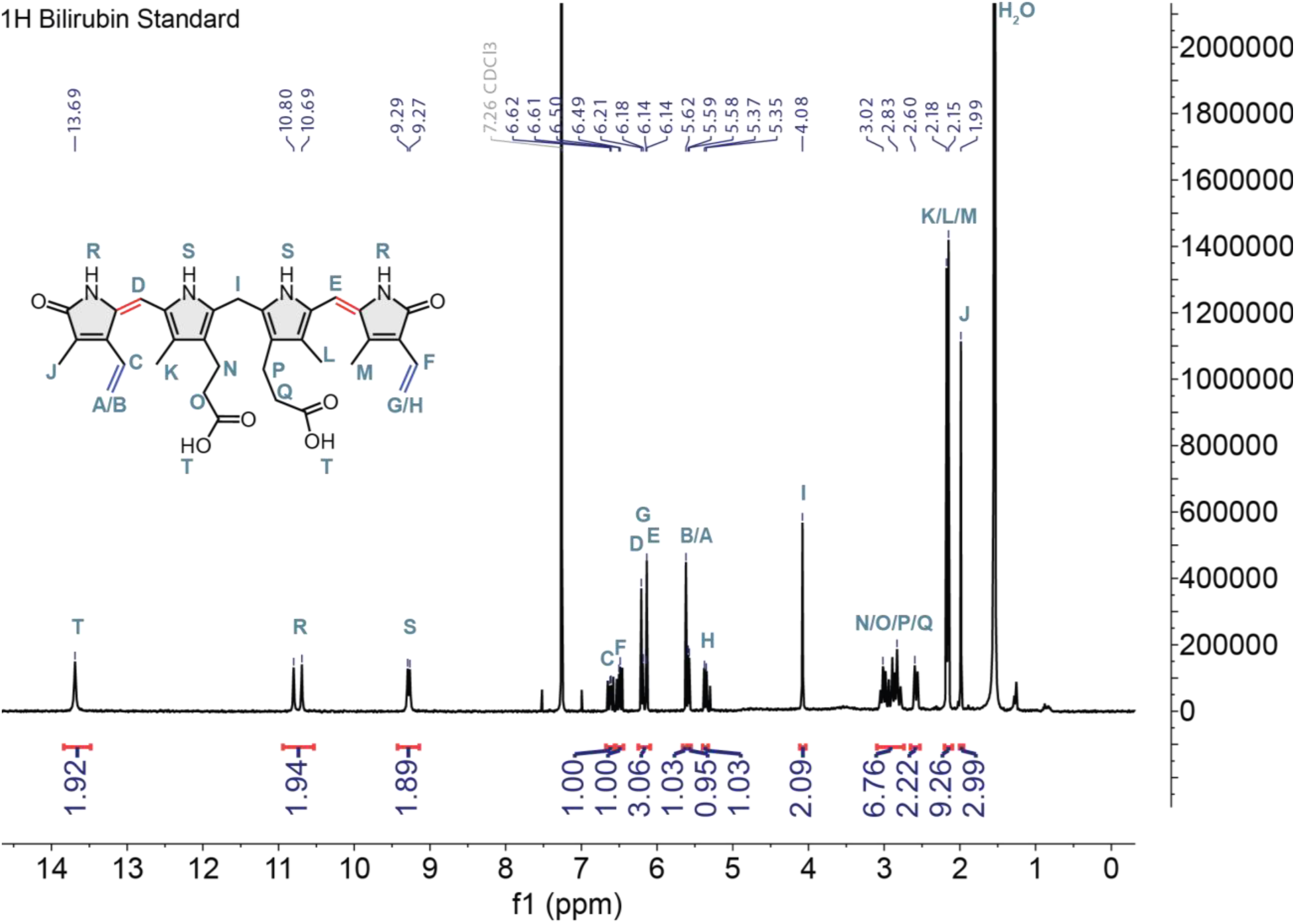
Full ^1^H NMR spectrum of commercially obtained bilirubin.

**Supplemental Figure 4:**
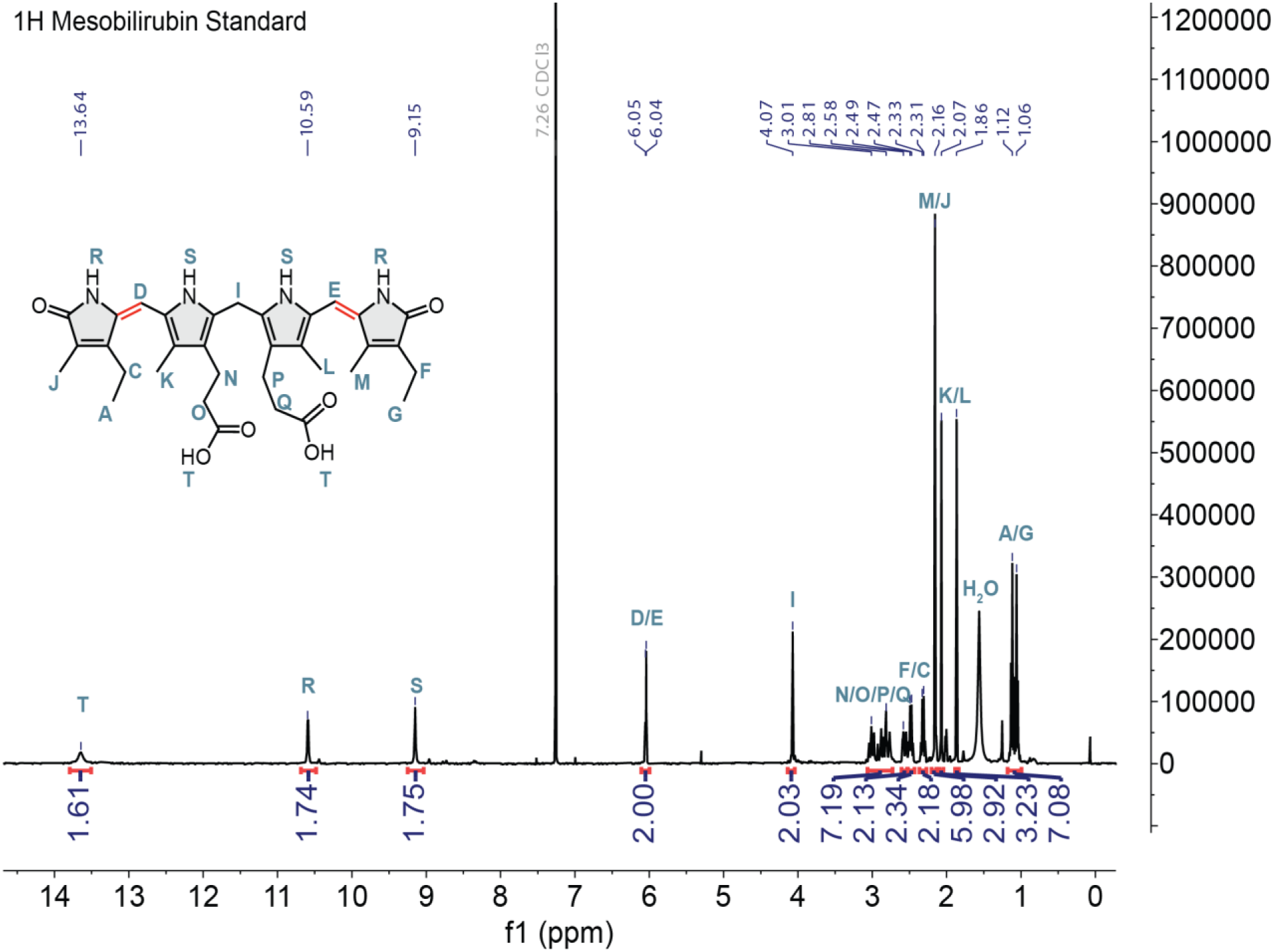
Full ^1^H NMR spectrum of commercially obtained mesobilirubin.

**Supplemental Figure 5:**
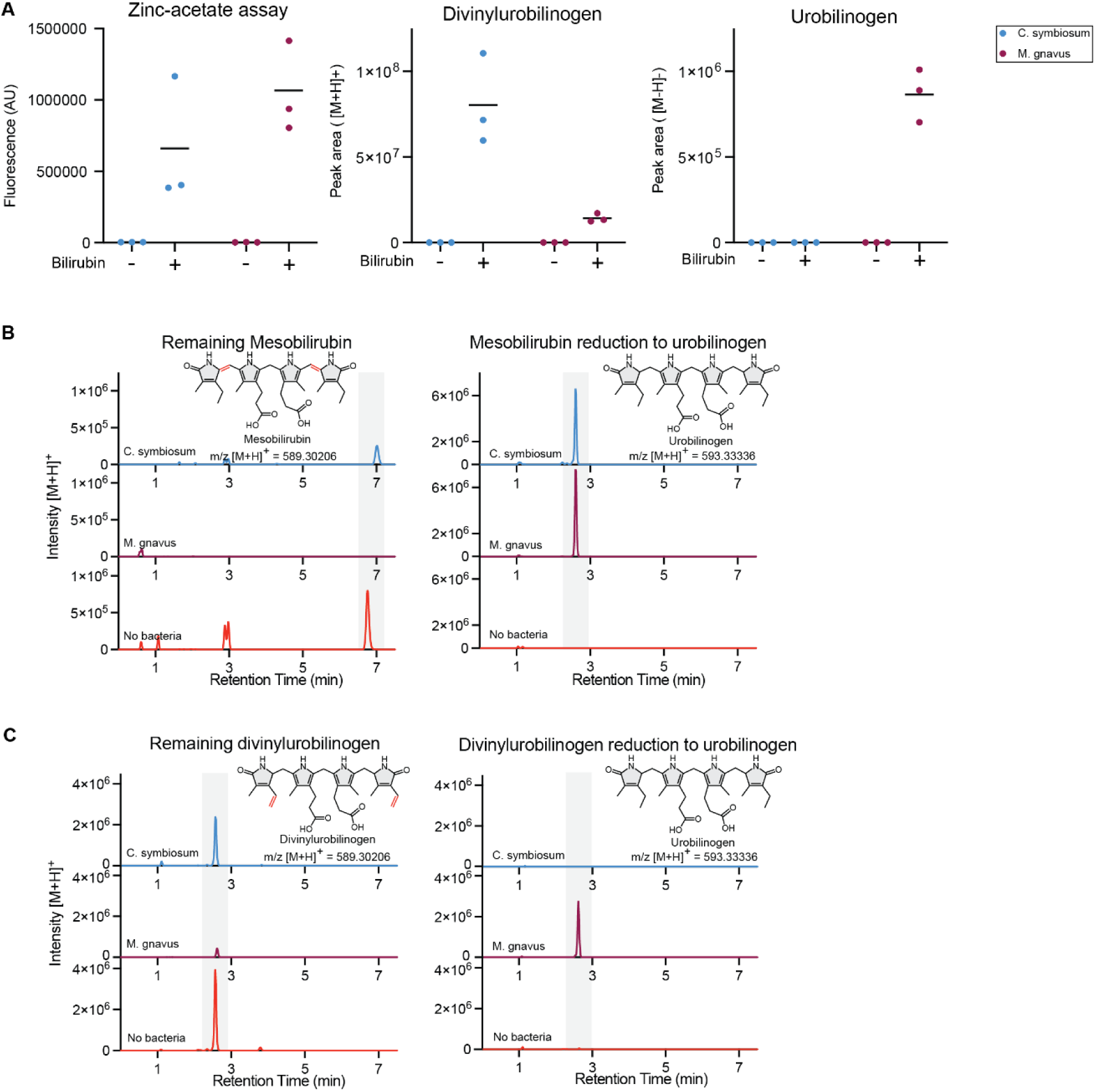
BilR-encoding strains can reduce methine groups of mesobilirubin. **A.** Fluorescence intensity at 495 nm wavelength excitation and 525 nm wavelength emission of crude extract of bacteria incubated with 10 μM bilirubin for 24 hours (left). Each dot represents average of three reads from one culture. Paired LC-MS detection of divinylurobilinogen (middle) and urobilinogen (right) from the same samples as processed in the zinc-acetate fluorescence assay. Data are represented as mean and individual values of replicate cultures. **B.** Extracted ion chromatogram for mesobilirubin (left, m/z=589.30) and urobilinogen (right, m/z=593.33) for *C. symbiosum* (blue), *M. gnavus* (purple), or media control (red) incubated with 1 μM mesobilirubin for 24 hours. **C.** Same as in B, but incubated with 10% spent media containing divinylurobilinogen.

**Supplemental Figure 6:**
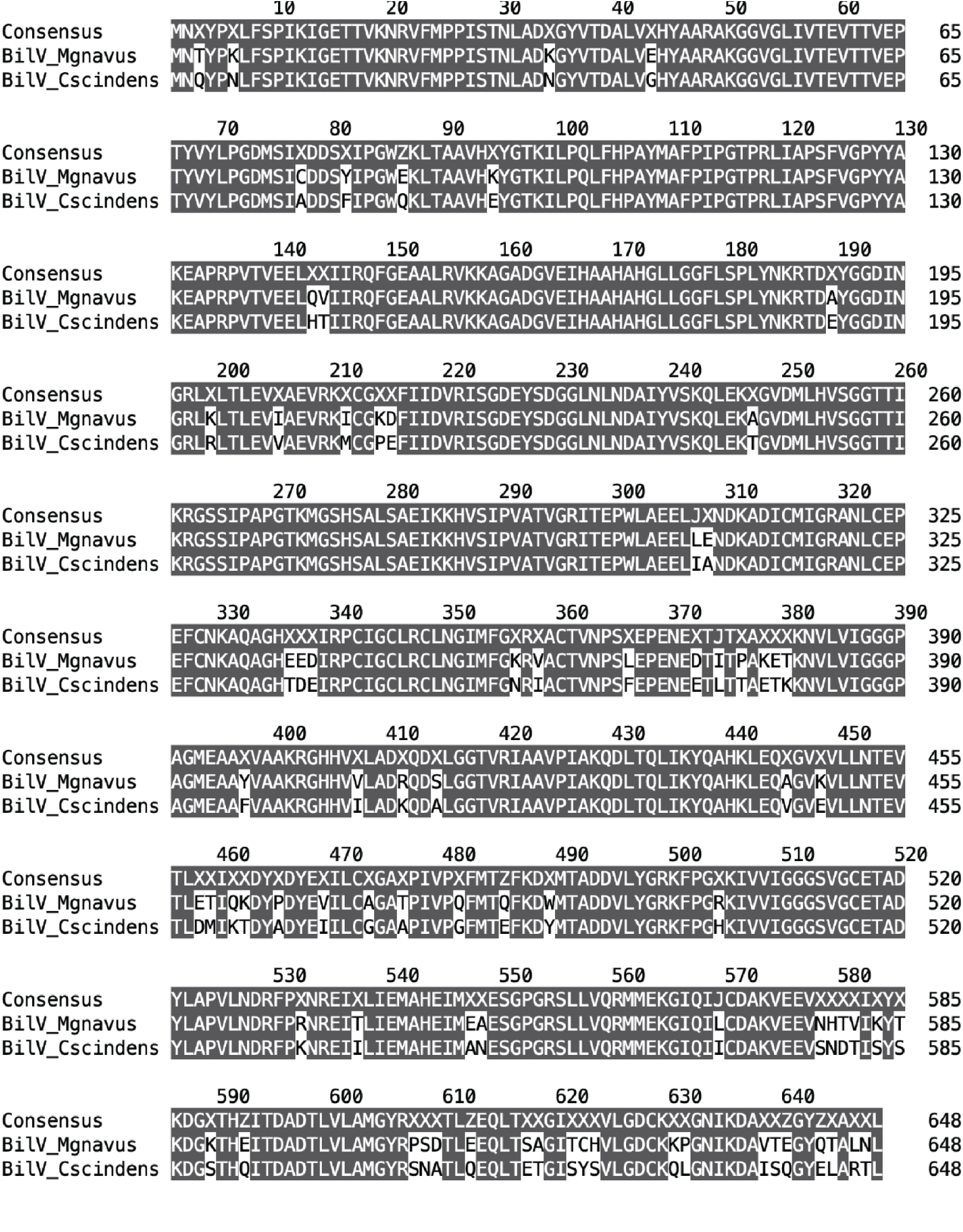
Alignment of bilinoid vinyl reductase from *M. gnavus* and *C. scindens*. Pairwise alignment of bilinoid vinyl reductases using Geneious Alignment global alignment with free end gaps (default parameters).

**Supplemental Figure 7:**
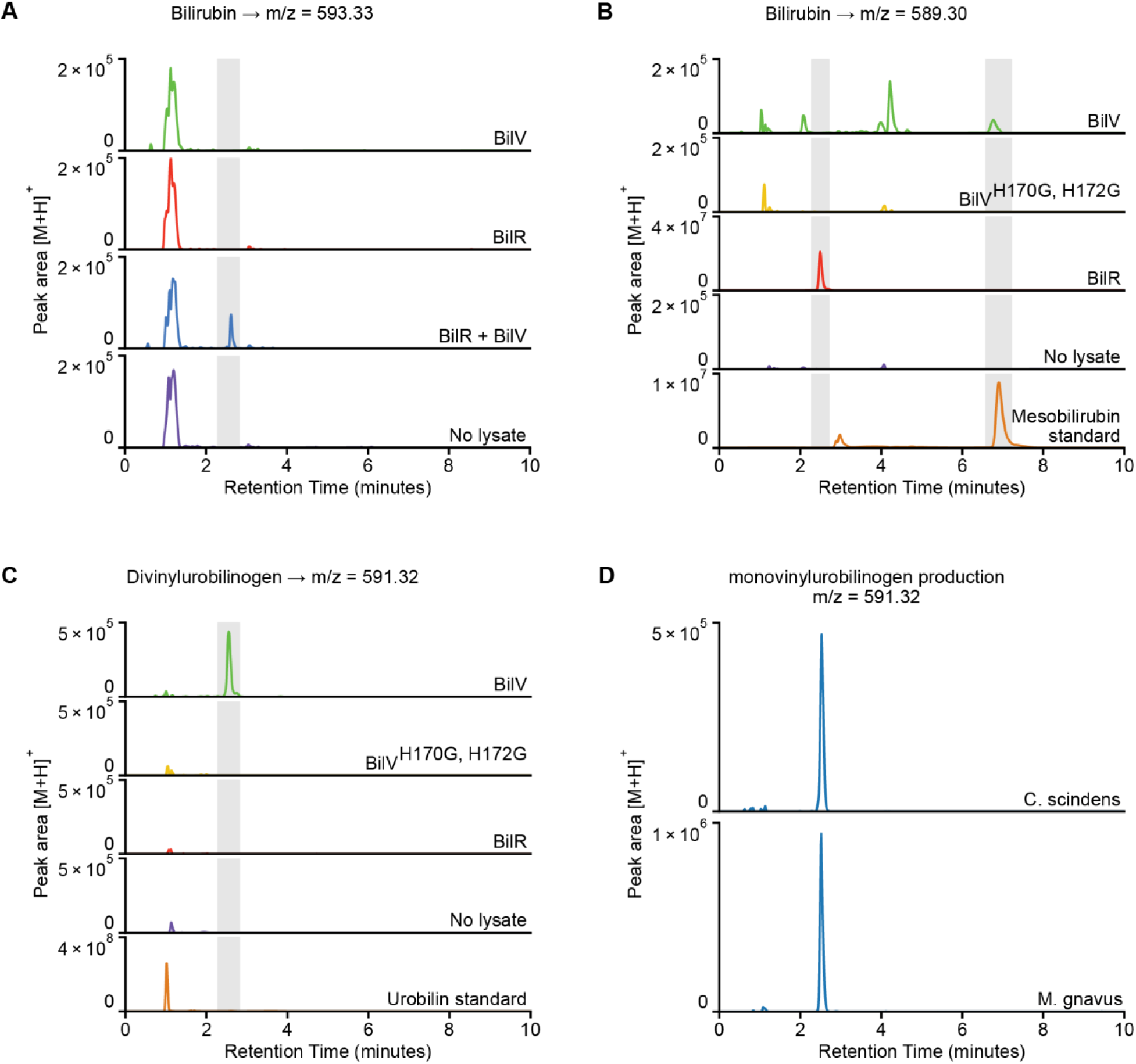
Representative extracted ion chromatograms from *E*. *coli* lysates expressing protein of interest to characterize activity of BilV. **A-C.** Representative extracted ion chromatograms for **A.** urobilinogen (m/z = 593.33, RT = 2.6 min), **B.** compounds with m/z= 589.30, consistent with divinylurobilinogen (RT = 2.5) and mesobilirubin (RT = 6.7), and **C.** compounds with m/z = 591.31, consistent with monovinylurobilinogen, for quantified plots presented in Figure 4A-D. **D.** Representative extracted ion chromatograms for compound with m/z = 591.32, consistent with monovinylurobilinogen, from *M*. *gnavus* incubated with bilirubin or *C*. *scindens* incubated with divinylurobilinogen.

**Supplemental Figure 8:**
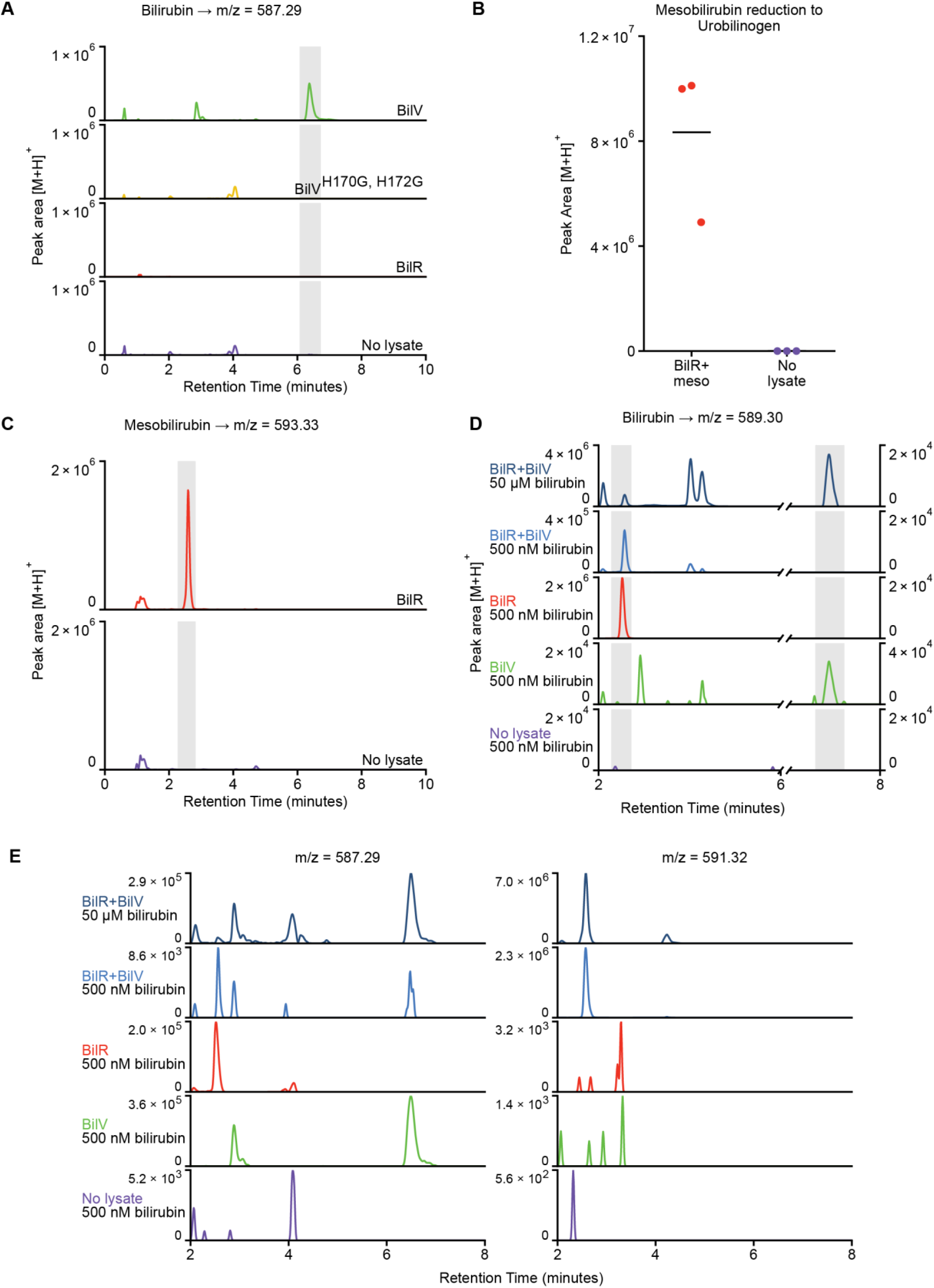
Representative extracted ion chromatograms from *E*. *coli* lysates expressing protein of interest to characterize architecture of the bilirubin metabolism pathway. **A.** Representative extracted ion chromatogram for compounds with m/z = 587.29, consistent with monovinylbilirubin, from lysates of *E*. *coli* expressing indicated proteins and incubated with 50 µM bilirubin for 6 hours, for quantified plot presented in Figure 4D. **B.** Quantified peak area and **C.** representative extracted ion chromatogram of urobilinogen (m/z=593.33, retention time=2.6 min) from lysates of *E. coli* expressing BilR and incubated with 500 nM mesobilirubin for 6 h.**D.** Representative extracted ion chromatograms of m/z values corresponding to compounds with m/z =589.30, consistent with divinylurobilinogen, mesobilirubin, or compounds with reduction of one vinyl group and one methine bridge, from lysates of *E*. *coli* expressing indicated proteins and incubated with bilirubin (500 nM, except where indicated otherwise) for 6 hours. **E.** Representative extracted ion chromatograms of m/z values corresponding to additional putative intermediates of bilirubin metabolism from lysates of *E*. *coli* expressing indicated proteins and incubated with bilirubin (500 nM, except where indicated otherwise) for 6 hours.

**Supplemental Figure 9:**
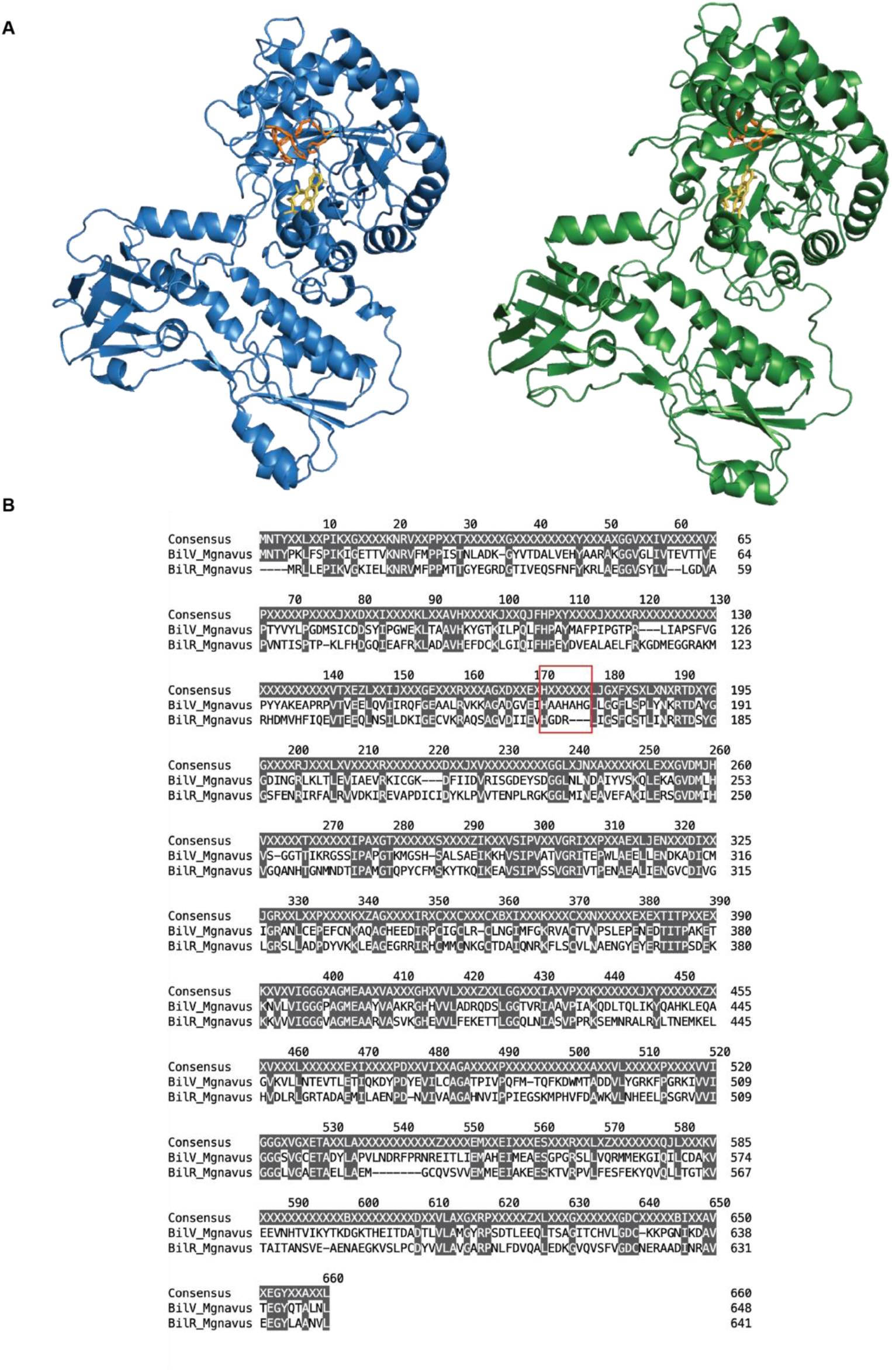
Comparison of bilinoid vinyl reductase and bilirubin reductase. **A.** Alphafold3-predicted models of bilirubin reductase (left) and bilinoid vinyl reductase (right) from *M. gnavus* modeled with flavin mononucleotide (highlighted in yellow). Proposed conserved residues are highlighted in orange; for bilirubin reductase HGDR164-168, for bilinoid vinyl reductase AHAH169-172. **B.** Pairwise alignment of bilinoid vinyl reductase and bilirubin reductase from *M. gnavus* using Geneious Alignment global alignment with free end gaps (default parameters). Red box highlights the conserved residues proposed to define substrate specificity.

**Supplemental Figure 10:**
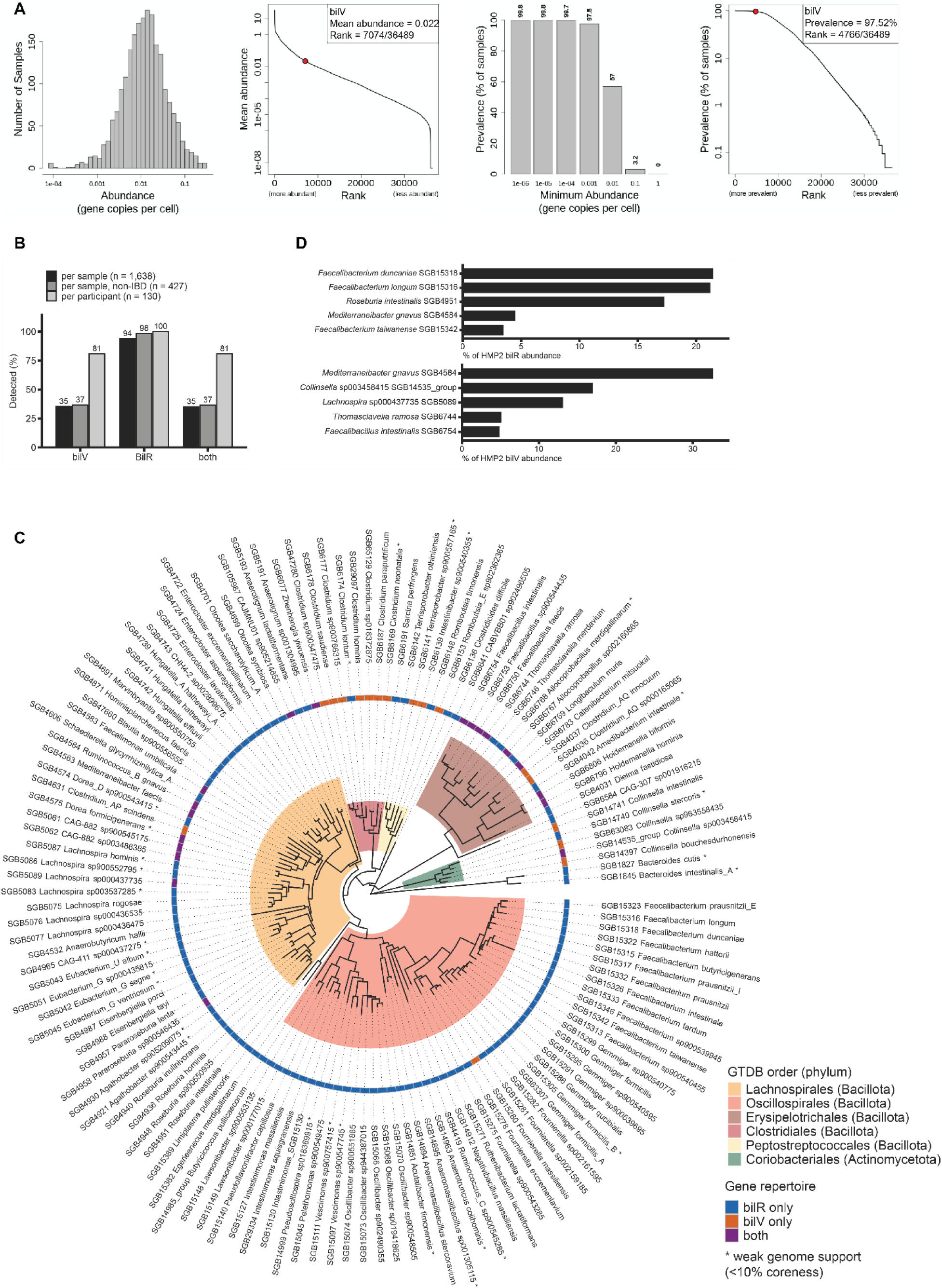
Bilinoid vinyl reductase is prevalent and abundant in the human microbiome. **A.** Abundance and prevalence of *bilV* in metagenomes of gut microbiomes of healthy humans, as determined with MetaQuery. **B.** Prevalence of *bilV* and *bilR* in metagenomes of gut microbiomes in the HMP2 cohort. **C.** Phylogenetic tree of species genome bins (SGBs) that encode *bilV* or *bilR* in metagenomes from the HMP2 dataset. Shading of clades corresponds to order. Weak genome support is defined as none of the gene families assigned to an SGB reaching 10% coreness in the pangenome and the SGB contributing less than half of the highest abundance for every assigned gene family. **D.** Contributions of individual SGBs to total abundance of *bilV* and *bilR* in metagenomes from the HMP2 dataset. Top 5 SGBs are shown for each.

